# Insights into plant-part specific N_2_O production in roots and shoots of chicory (*C. intybus*) using stable isotope labelling

**DOI:** 10.1101/2025.10.28.685070

**Authors:** Moritz Schroll, Maurice Maas, Steffen Greiner, Thomas Klintzsch, Frank Keppler

## Abstract

- Nitrous oxide (N_2_O) substantially contributes to climate change and stratospheric ozone degradation, yet large uncertainties in its global budget indicate unknown or overlooked sources. Increasing evidence suggests that plants may also produce N_2_O, though the underlying mechanisms and pathways remain poorly constrained.
- To examine whether plants can form N_2_O under sterile conditions and to assess the contribution of different plant parts, we applied a novel ^15^N stable isotope labelling approach using sterile *Cichorium intybus* root and shoot cultures incubated separately under light and dark conditions.
- All root/shoot cultures showed N_2_O formation under dark conditions, whereas shoots under light showed reduced or even uptake of N_2_O, indicating photosynthetically driven suppression of formation pathways or simultaneous internal degradation of N_2_O.
- Isotopic analyses revealed distinct formation pathways: roots supplemented with ^15^NO_3_⁻ showed position-specific ^15^N enrichment consistent with N_2_O formation via nitric oxide as an intermediate, linking root-derived N_2_O to NO_3_⁻ reduction. In contrast, root/shoot incubations with ^15^N-NH_4_^+^ supplementation and shoots in darkness emitted N_2_O without clear ^15^N enrichment suggesting alternative formation pathways independent of these compounds. Our isotopic labelling approach powerfully disentangled N_2_O formation mechanisms yet highlights necessary further exploration of plant N_2_O cycling to improve global budgets and enable potential mitigation strategies.

## Introduction

Nitrous oxide (N_2_O) is the third most important anthropogenic greenhouse gas, contributing to climate forcing and likely emerging as the dominant ozone-depleting substance emitted during the 21^st^ century (Forster *et al*., 2007; Ravishankara *et al*., 2009). Its atmospheric concentration has increased by approximately 22% compared to preindustrial levels, underscoring its growing impact on global climate and stratospheric ozone depletion (IPCC, 2014; Tian *et al*., 2020). Nitrogen, a vital macronutrient, plays a crucial role in plant growth and development. It is a fundamental component of chlorophyll, amino acids, nucleic acids, and other secondary metabolites. Plants primarily acquire nitrogen in the forms of nitrate (NO_3_^−^) and ammonium (NH_4_^+^) through processes like nitrification and ammonification in the natural nitrogen cycle, as well as through fertilizers (Zayed *et al*., 2023). NO_3_^−^ and NH_4_^+^, particularly in soils are further converted to N_2_O by microorganisms through nitrification, nitrifier-denitrification and denitrification processes and are responsible for two thirds of N_2_O emissions (Tian *et al*., 2020). Traditionally, studies on N_2_O cycling in terrestrial ecosystems have focused on the exchange of N_2_O between soil surfaces and the atmosphere. While plants have primarily been recognized as conduits for N_2_O transfer in the soil-plant-atmosphere system, their own specific role in N_2_O formation and involved processes have surprisingly only recently been more in the focus of scientific interest. Already in the 1980s Dean & Harper (1986) described the formation of N_2_O from two different plants species. Since then, an increasing number of studies observed the formation of N_2_O production and subsequent emission from vegetation including trees, plants, shrubs, herbs, cryptogamic covers and grass (Dean & Harper, 1986; Chen *et al*., 1999; Goshima *et al*., 1999; Smart & Bloom, 2001; Hakata *et al*., 2003; Pihlatie *et al*., 2005; Bruhn *et al*., 2014; Lenhart *et al*., 2015, 2019; Machacova *et al*., 2016). However, there are contrasting findings, with several studies reporting N_2_O formation from the plants themselves, for example, when grown under sterile conditions (e.g., Lenhart *et al*., 2019) opposed to N_2_O production in plants attributed mainly to endophytic and phyllospheric bacteria (Bowatte *et al*., 2015; Qin *et al*., 2024). This contrasting evidence clearly shows that, even almost 40 years after its first description, there is a clear lack of understanding of the underlying processes linked with plant N_2_O formation. This is further highlighted by the reported N_2_O emission rates from plants showing a large variability ranging from 10 to 6000 pmol g^-1^ h^-1^ (Lenhart *et al*., 2019; Qin *et al*., 2024). Nevertheless, recent studies by Lenhart *et al*. (2019) and Qin *et al*. (2024) estimate that plants contribute between 6% and 11% of global annual N_2_O emissions. These findings suggest that plant-mediated N_2_O emissions are a ubiquitous process and highlight their substantial role in the global N_2_O cycle. Despite this, terrestrial vegetation has not been considered in global N_2_O budgets (Lenhart *et al*., 2019).

However, even though vegetation has been identified as a potentially large contributor to global N_2_O emissions, the mechanisms of N_2_O formation are mostly unclear. Previous studies have started to provide insight into the N_2_O formation mechanism using stable isotope techniques, since it allows for the differentiation of multiple sources due to their individual isotopic fingerprint. Smart & Bloom (2001), by employing ^15^N-labelled NO_3_^−^ as a tracer, demonstrated that enzymatic processes in plant foliage can generate N_2_O. In a related study, Hakata *et al*. (2003) observed the emission of ^15^N-labelled N_2_O from 16 different plant taxa grown aseptically in a ^15^N-labelled NO_3_^−^ medium, concluding that NO_3_^−^ conversion to N_2_O is widespread in plants. Lenhart *et al*. (2019) were the first to report isotopic signatures of N_2_O released by plants, in addition to measuring the emission rates of 32 plant species from 22 families under controlled laboratory conditions. Emerging research suggests that plants possess the ability to denitrify (Timilsina *et al*., 2020, 2022a). For instance, mitochondria may serve as sites for N_2_O formation in plant cells, with evidence showing that plant N_2_O fluxes correlate positively with respiration rates and negatively with surface areas (Timilsina *et al*., 2022a). These relationships suggest that plant roots and mitochondria are integral to N_2_O production in natural conditions. Furthermore, NO reduction to N_2_O through the phytoglobin-NO cycle may represent an important mechanism for reducing nitro-oxidative stress, enhancing plant survival under hypoxic conditions (Timilsina *et al*., 2022a). Experimental studies have also shown that NO_3_^−^ is a critical precursor compound for plant-mediated N_2_O production. NO_3_^−^, undergoes reduction and assimilation in leaves during photosynthesis, ultimately producing N_2_O (Dean & Harper, 1986; Goshima *et al*., 1999; Smart & Bloom, 2001; Hakata *et al*., 2003). The reductive pathway in mitochondria further reduces NO, formed under low oxygen (O_2_) conditions to N_2_O. This mechanism not only mitigates NO toxicity but may also represent a significant biogenic source of N_2_O globally (Timilsina *et al*., 2020). Beyond microbial N_2_O formation, abiotic mechanisms, such as UV-induced N_2_O release from plant surfaces, also contribute to N_2_O fluxes (Bruhn et al., 2014). Building on existing laboratory studies clearly identifying N_2_O formation from plants, two recent studies for the first time addressed the measurement of plant related N_2_O formation in the field and disentangling this production pathway from soil N_2_O fluxes using the stable isotope composition of N_2_O (Timilsina *et al*., 2022b; Xia *et al*., 2024). By comparing the isotopic signatures and emissions of N_2_O released from plants and soils, these studies unambiguously showed that plants are not merely passive conduits for N_2_O produced in soils, but are themselves active sources contributing to atmospheric N_2_O fluxes. Moreover, there is further emerging evidence, that N_2_O is not only produced by plants, but that plants are also capable of reducing N_2_O fluxes, especially under photosynthetic activity (Schützenmeister *et al*., 2020). This finding has led to the hypothesis that N_2_O formation in plants may be more locally linked with the root systems (Timilsina *et al*., 2022b).

However, currently no study has systematically investigated the difference between N_2_O formation in different organs of plants such as the root and shoot. Thus, in this study, we investigated the N_2_O emissions and isotopic composition of N_2_O from incubated chicory shoot and root cultures cultivated under sterile conditions using stable isotope labelling experiments with ^15^N-NO_3_^−^ and ^15^N-NH_4_^+^ under different light conditions.

## Materials and Methods

### Cultivation of root and shoot cultures of *Cichorium intybus*

Chicory (*Cichorium intybus*) was selected for this study due to its physical traits, which allow for the separate cultivation of root and shoot cultures, as well as considerable in-house practical experience in the sterile cultivation. This allowed us to differentiate between N_2_O formation taking place in these different plant organs without potential interference of the other plant part. Chicory root cultures were established and cultivated following the methodology described by Kusch *et al*., 2009. Root cultures were maintained in Erlenmeyer flasks filled with liquid medium and cultivated at a temperature of 25 °C in a shaking incubator at 90 rpm in darkness. For Chicory shoot cultures, chicory seeds were first treated with sodium hypochlorite and 0.01% Triton X-100, followed by submersion in ethanol and subsequent rinsing with sterile water. Seeds were then germinated on Gamborg B5 medium (Duchefa Biochemie, Netherlands) supplemented with 1% sucrose and 0.8% plant agar (Duchefa Biochemie) with a light phase (16 h) and a dark phase (8 h). 6 weeks after germination, the seedlings were cut and the shoot segments were separately cultivated.

### Experimental setup and conditions

The experimental setup of this study consisted of separately incubated chicory shoots and roots with the addition of ^15^N-labeled NH_4_^+^ and NO_3_^−^ solutions. Stable isotope labelled solutions were prepared by dissolving ^15^N-labelled ammonium chloride (NH_4_Cl, ^15^N ≥ 98%, Sigma-Aldrich, USA) and sodium nitrate (NaNO_3_, ^15^N ≥ 98%, Sigma-Aldrich, USA) in demineralised and autoclaved water. The final concentrations of NH_4_Cl and NaNO_3_ ranged between 27 and 34 mmol L^-1^. All incubated roots and shoot cultures as well as the respective controls were incubated in airtight 580 mL glass flasks (Weck, Germany) equipped with silicone stoppers (Saint-Gobain Performance Plastics, France) for headspace sampling. Root cultures were incubated in liquid Gamborg B5 medium at 23 °C and light conditions. Since roots are not capable of photosynthesis, light exposure was not expected to influence this experiment. Accordingly, root controls consisted of the same liquid medium and glass flasks that were used for root cultures, but without the presence of the roots. Chicory shoots were incubated with a Gamborg B5 medium under both dark and light conditions to assess potential effects of light on N_2_O formation and isotopic composition. Control experiments consisted of the same medium without shoot cultures. A volume of 10 mL of the prepared ^15^N-labelled solutions were added to the shoot and root cultures as well as controls using a laminar flow bench. Additionally, shoots and controls were incubated in the dark without the addition of ^15^N-NH_4_^+^ or ^15^N-NO_3_^−^ solutions (Table 1). Due to technical difficulties no root and ‘shoot + light’ without added labelled compounds could be investigated. Following the addition of the labelled solutions, the glass flasks were transferred to a climate chamber with a temperature of 25 °C. Dark experiments were carried out with the chamber lights switched off, whereas light conditions followed a day-night cycle with a 16 h light phase and an 8 h dark phase. The intensity of photosynthetically active radiation was ∼90 μmol s^-1^ m^-2^.

**Table 1:** Overview of incubation setup, added ^15^N-labelled NH_4_^+^ or NO_3_^−^ solutions, light conditions and the number of replicates (n).

| Plant part | Treatment | Light condition | n |
| --- | --- | --- | --- |
| Root culture | <sup>15</sup>N-NO<sub>3</sub><sup>-</sup> | dark | 3/4 |
|  | <sup>15</sup>N-NH<sub>4</sub><sup>+</sup> |  |  |
| Control | <sup>15</sup>N-NO<sub>3</sub><sup>-</sup> | dark | 2 |
|  | <sup>15</sup>N-NH<sub>4</sub><sup>+</sup> |  |  |
| Shoot culture | <sup>15</sup>N-NO<sub>3</sub><sup>-</sup> | light | 2/4 |
|  | <sup>15</sup>N-NH<sub>4</sub><sup>+</sup> |  | 3/4 |
|  | <sup>15</sup>N-NO<sub>3</sub><sup>-</sup> | dark | 3/4 |
|  | <sup>15</sup>N-NH<sub>4</sub><sup>+</sup> |  | 4 |
|  | none |  | 3/4 |
| Control | <sup>15</sup>N-NO<sub>3</sub><sup>-</sup> | light | 3 |
|  | <sup>15</sup>N-NH<sub>4</sub><sup>+</sup> |  |  |
|  | <sup>15</sup>N-NO<sub>3</sub><sup>-</sup> | dark | 4 |
|  | <sup>15</sup>N-NH<sub>4</sub><sup>+</sup> |  |  |
|  | none |  |  |

### Gas sampling

First the internal pressure inside the glass flasks was measured using a hand-held manometer (GHM Messtechnik GmbH, Germany) to allow the calculation of the exact amount of substance of target gases. Subsequently, for the analysis of N_2_O mixing ratios and its isotopic composition a plastic syringe (Plastipak, USA) was used to extract 60 mL of sample gas which was transferred into 20 mL or 40 mL glass vials equipped with silicone-PTFE septa or butyl-PTFE septa. For the analysis of oxygen (O_2_) and carbon dioxide (CO_2_) mixing ratios, a 7 mL gas sample was injected into 3 mL exetainers (Labco Limited, UK). To compensate for the changing pressure inside the glass flasks resulting from gas sampling, a cannula equipped with a 0.2 µm filter (Cytiva, USA) was used. Gas samples were collected at three different time points: 6, 24 and 48 hours after start of the incubation. In addition, four laboratory air samples were taken at the start of the experiment to determine the initial gas composition within the glass flasks.

### Isotopes of N_2_O and definition of *δ*-values

The four most common isotopologues of nitrous oxide (N_2_O) are ^14^N^14^N^16^O, ^14^N^14^N^18^O, ^14^N^15^N^16^O, and ^15^N^14^N^16^O. The isotopomers ^14^N^15^N^16^O and ^15^N^14^N^16^O differ only in the position of the ^15^N atom within the molecule. When the ^15^N atom is located at the central nitrogen position, this configuration is referred to as ^15^N^α^, whereas its placement at the terminal position is defined as ^15^N^β^. The mean of the isotopic ratios of ^15^N^α^ and ^15^N^β^ is known as ^15^N^bulk^ (Toyoda & Yoshida, 1999). In this study, all stable nitrogen isotope ratios are expressed using the conventional “delta” (δ) notation (Eq.1), referring to the relative difference of the isotope ratio of a sample from the *δ*^15^N_air_ standard:

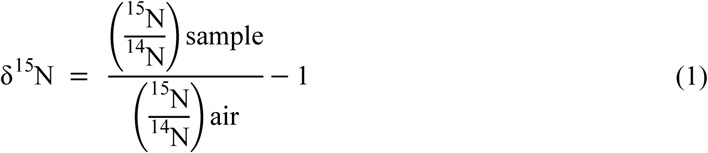

We support the proposal of Brand & Coplen (2012) to use the term “urey” (Ur) as the unit for the isotopic delta in accordance with the guidelines of the International System of Units (SI). Therefore, in the following chapters, the relationship applies: 1 mUr = 1 ‰.

### Analysis of N_2_O mixing ratios and δ^15^N-N_2_O values

Analyses of N_2_O mixing ratios and its stable isotopes were performed using a Picarro G5131-*i* cavity ring-down spectroscope (CRDS, Picarro, USA). A Small Sample Introduction Module 2 (SSIM2, Picarro, USA) was used to analyse small sample volumes ranging from 20-50 mL. Synthetic air (20,5% O_2_ in N_2_, ALPHAGAZ™ 1, Air Liquide, France) was used as carrier gas. Between each run, the system was evacuated and flushed with synthetic air to maintain accuracy. To ensure high quality analysis, two standard gases with known mixing ratios and isotopic composition were measured at the beginning and end of each measurement day for calibration and drift correction. CRDS instruments are known to be affected by cross-sensitivities, where the occurrence and concentration of other gases can substantially influence the results of the analysis of the target gas (e.g., Harris *et al*., 2020). To prevent cross-sensitivities with volatile organic compounds, a glass tube filled with Tenax© (Sigma-Aldrich, USA) was placed in front of the inlet valve of the SSIM2 for the adsorption of volatile organic compounds. Additionally, the plastic syringe used for gas injection was partially filled the CO_2_ adsorbent Ascarite(II)© (Acros Organics, Belgium). Furthermore, corrections were applied to account for the influences of cross-sensitivities for CH_4_, O_2_, and CO_2_, the concentration of N_2_O, as well as for the effects of used glass vials, septa, and storage time. These corrections, conducted similar to Harris *et al*. (2020) are described in detail in the supplementary Fig. S1 and Method S1-3.

### Analysis of CO_2_ and O_2_ mixing ratios

CO_2_ and O_2_ mixing ratios were determined using a GC-2010 Plus gas chromatograph (Shimadzu, Japan) coupled with a barrier discharge ionization detector (BID-2010 Plus, Shimadzu, Japan). The setup included a packed stainless-steel column with a length of 2 m, an inner diameter of 0.53 mm and a carbon molecular sieve (ShinCarbon ST, 80/100 Mesh; Supelco, USA) as the stationary phase. High-purity helium (grade 6.0, Air Liquide, France) was used as the carrier gas at a constant flow rate of 5 mL min^-1^. Sample injection was performed using an autosampler (AOC-20i, Shimadzu, Japan). As part of quality control, one reference standard was measured after every six to nine single measurements. Calibration was conducted using certified standard gas mixtures (CRYSTAL Gas Mixture, Air Liquide, France). The quantification of CO_2_ and O_2_ was determined by analysing multiple reference standards (CO_2_: 400 ppm–40% by volume, O_2_: 1–22% by volume), with each measurement performed in triplicate.

### Statistical methods

The evaluation of measurement data from the Picarro G5131-*i* was performed using R and Microsoft Excel. For the analysis of raw data from the Picarro G5131-*i*, moving averages of the measurements over 100 seconds were calculated. For each measurement, a plateau period of at least 5 minutes was selected. Mixing ratios, production rates and the isotopic change rate are presented as the arithmetic mean (*n* = 3 or 4) of independent replicates with their standard deviations (SDs). For some samples, results at specific time points were discarded due to analytical problems. If fewer than three measurements were available for a given time point, no calculation of the SD was carried out. Additionally, the obtained data were used to determine the production rates of the gases N_2_O, O_2_, and CO_2_ for the shoot and root cultures as well as for the controls. For each incubated glass flask, an individual production rate was determined using linear regression. The arithmetic means of these production rates were then calculated for the same plant compartment and treatment. Furthermore, t-tests were performed using SigmaPlot (v.12.2.0.45, USA) to statistically support the observed findings. Please note, that his study follows the recommendations of the American Statistical Association, ensuring that p-values and other statistical parameters were not used as the sole basis for drawing conclusions (Wasserstein *et al*., 2019). Consequently, the term “statistically significant” was deliberately avoided in the interpretation of the results.

## Results

The incubations comprised a series of experiments in which shoot and root cultures of chicory were incubated under dark and light conditions with the addition of ^15^N-labeled NO_3_^−^ and NH_4_^+^ to assess the changes in the mixing ratios of N_2_O, CO_2_ and O_2_ as well as the isotopic composition of N_2_O. Please note that the results are presented as changes relative to the measurements at the start of the incubation. Furthermore, measured values are reported as absolute changes in the respective glass flasks, without considering the biomass of the different plant parts to ensure the comparability of flasks with plants and the respective controls, as the controls do not possess biomass. When calculating production rates and isotopic change rates dry plant biomass was considered.

### Chicory roots under dark conditions

N_2_O mixing ratios during the incubation of chicory roots increased by 38.8 ± 4.6 ppbv when supplemented with ^15^NO_3_^−^ (*n* = 3, Fig. 1a). The increase in N_2_O mixing ratios was accompanied by a strong shift in stable isotope values towards more positive values with δ^15^N^bulk^-N_2_O values increasing by 64.7 ± 6.5 mUr (*n* = 3), δ^15^N^α^-N_2_O by 72.0 ± 7.6 mUr (*n* = 3) and δ^15^N^β^-N_2_O values 57.7 ± 5.4 mUr (*n* = 3) (Fig. 1d, e, f). CO_2_ and O_2_ mixing ratios exhibited opposite trends, with CO_2_ increasing on average by 20.4 ± 0.7% (*n* = 3) while O_2_ mixing ratios decreased by 15.0 ± 1.1% (*n* = 3) (Fig. 1b, c). When chicory roots were supplemented with ^15^NH_4_^+^, N_2_O mixing ratios increased by 46.5 ± 8.4 ppbv (*n* = 3; Fig. 1a). Compared to the treatment with NO_3_^−^, ‘root + ^15^NH_4_^+^’ showed a less pronounced shift towards more positive stable isotope values with δ^15^N^bulk^-N_2_O values increasing by 13.3 ± 1.3 mUr (*n* = 3), δ^15^N^α^-N_2_O values by 17.1 ± 0.7 mUr (*n* = 3) and δ^15^N^β^-N_2_O values by 9.6 ± 1.9 mUr (*n* = 3) (Fig. 1d, e, f). During the incubation period, CO_2_ mixing ratios increased by 21.9 ± 1.2% (*n* = 3) while O_2_ mixing ratios decreased by 16.7 ± 1.0% (*n* = 3).

**Fig. 1:**
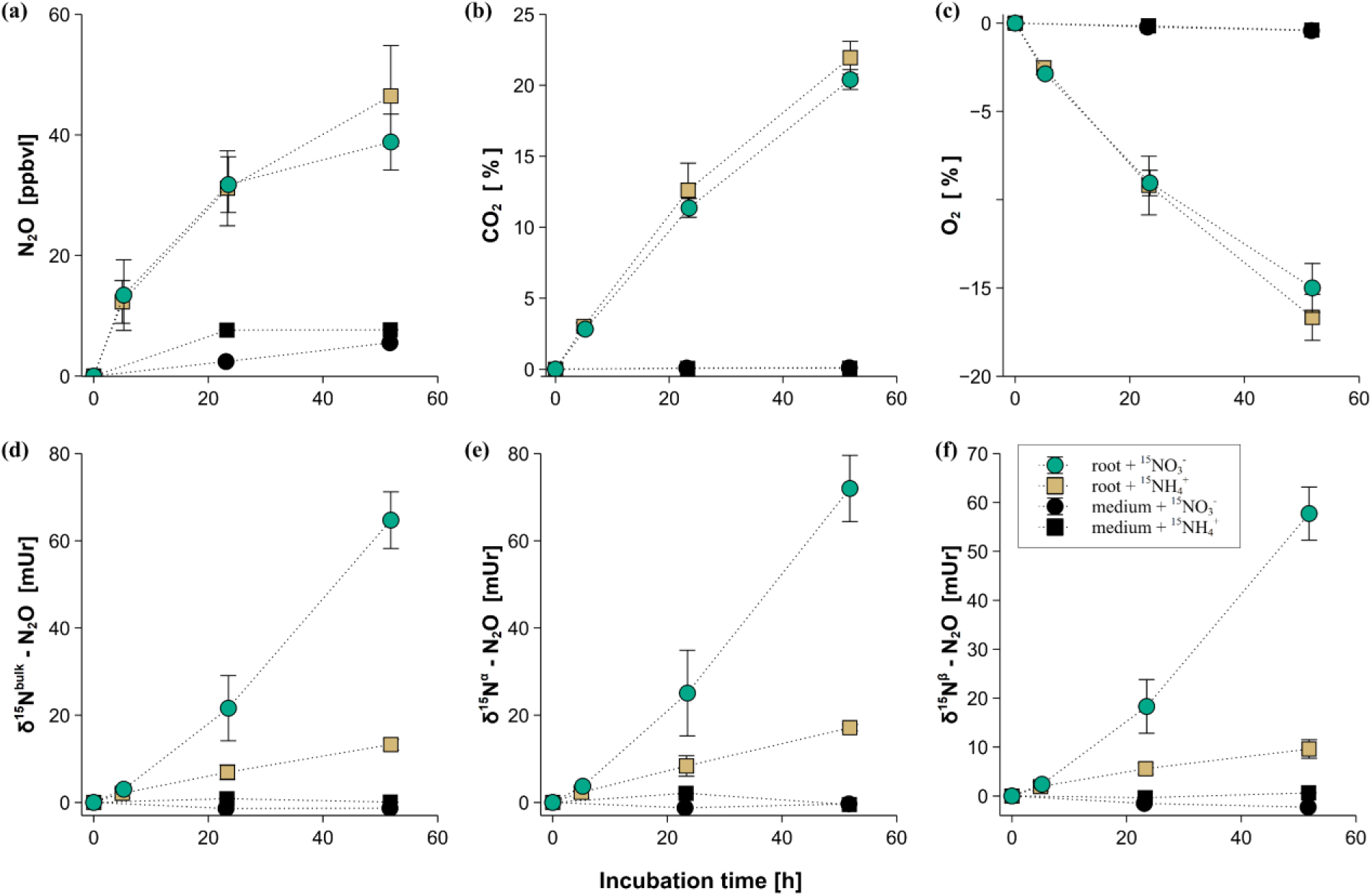
Relative changes of (a) N_2_O, (b) CO_2_ and (c) O_2_ mixing ratios and (d) δ^15^N^bulk^-, (e) δ^15^N^α^- and (f) δ^15^N^β^-N_2_O values during the dark incubation of chicory root cultures with supplementation of ^15^NO_3_^−^ and ^15^NH_4_^+^. Please note that the results are presented as changes relative to the measurements at the incubation start. Error bars represent the standard deviation (*n* = 3 or 4). Reported values reflect absolute changes in the respective glass flasks and were not normalized to biomass or its variation over time.

Controls containing ‘medium + ^15^NO_3_^−^’ and ‘medium + ^15^NH_4_^+^‘ showed only minor increases in N_2_O mixing ratios (5.5 ppbv for ‘medium + ^15^NO_3_^−^’ (*n* = 2) and 4.5 ppbv for ‘medium + ^15^NH_4_^+^’ (*n* = 2), respectively) and negligible changes in stable isotope values as well as CO_2_ and O_2_ mixing ratios during the incubation.

### Chicory shoots under dark conditions

N_2_O mixing ratios increased by 9.8 ± 2.8 ppbv when chicory shoots were incubated in the dark with ^15^NO_3_⁻ (*n* = 3; Fig. 2a). This increase was accompanied by pronounced shift towards more positive δ^15^N values: +10.1 ± 1.4 mUr for δ^15^N^bulk^–N_2_O, +15.8 ± 1.1 mUr for δ^15^N^α^–N_2_O and +4.6 ± 2.5 mUr for δ^15^N^β^–N_2_O (*n* = 3; Fig. 2b–d). CO_2_ mixing ratios increased by 3.28 ± 0.77 % while O_2_ decreased by 3.91 ± 0.59 % (*n* = 3; Fig. 2e,f).

**Fig. 2:**
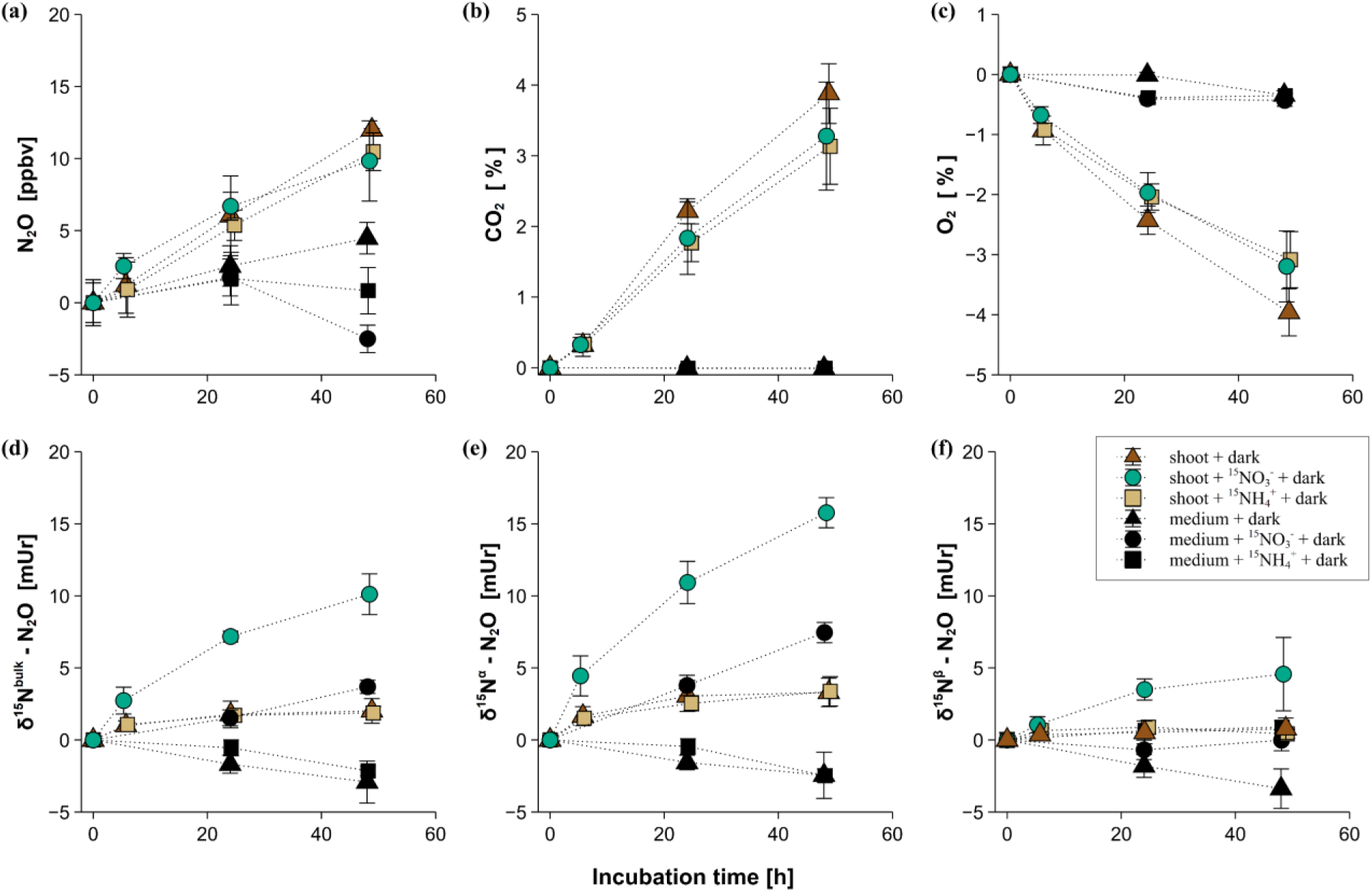
Relative changes of (a) N_2_O, (b) CO_2_ and (c) O_2_ mixing ratios and (d) δ^15^N^bulk^-, (e) δ^15^N^α^- and (f) δ^15^N^β^-N_2_O values during the incubation of chicory shoots in dark conditions with supplementation of ^15^NO_3_^−^ and ^15^NH_4_^+^. Please note that the results are presented as changes relative to the measurements at the incubation start. Error bars represent the standard deviation (*n* = 3 or 4). Reported values reflect absolute changes in the respective glass flasks and were not normalized to biomass or its variation over time.

When shoots were incubated in the dark with ^15^NH_4_⁺, N_2_O mixing ratios elevated by 10.5 ± 1.3 ppbv (*n* = 4; Fig. 2a). Isotopic shifts were smaller accounting for +1.9 ± 0.5 mUr for δ^15^N^bulk^–N_2_O, +3.4 ± 1.0 mUr for δ^15^N^α^–N_2_O and +0.4 ± 0.3 mUr for δ^15^N^β^–N_2_O (*n* = 4; Fig. 2b–d). CO_2_ increased by 3.13 ± 0.54 % and O_2_ decreased by 3.08 ± 0.47 % (*n* = 4; Fig. 2e,f).

In the ‘shoot + dark only’ treatment N_2_O mixing ratios rose by 12.0 ± 0.1 ppbv (*n* = 3), with δ^15^N^bulk^–N_2_O values increasing by +2.0 ± 0.9 mUr, δ^15^N^α^–N_2_O by +3.3 ± 1.0 mUr and δ^15^N^β^–N_2_O by +0.8 ± 0.7 mUr (*n* = 3). CO_2_ mixing ratios increased by 3.88 ± 0.42 % and O_2_ mixing ratios decreased by 3.96 ± 0.39 % (*n* = 3; Fig. 2e,f).

Control incubations (‘medium + dark + ^15^NO_3_⁻’; ‘medium + dark + ^15^NH_4_⁺’; ‘medium + dark only’) showed minimal N_2_O changes (–2.5 ± 1.0 ppbv, +0.8 ± 1.6 ppbv, and +4.5 ± 1.1 ppbv, respectively; *n* = 4) and negligible shifts in δ^15^N^bulk^–N_2_O values (+3.7 ± 0.5 mUr; –2.5 ± 0.5 mUr; –2.9 ± 1.5 mUr) for δ^15^N^α^–N_2_O/δ^15^N^β^–N_2_O respectively and CO_2_/O_2_ mixing ratios (< ± 0.4 %; *n* = 4).

### Chicory shoots under light conditions

The N_2_O mixing ratio increased by 15.4 ppbv when shoots were incubated in the light with ^15^NO_3_⁻ (*n* = 2; Fig. 3a). This was accompanied by a pronounced shift towards more positive δ^15^N values. δ^15^N^bulk^-N_2_O values increased by 14.4 mUr (*n* = 2; Fig. 3d), δ^15^N^α^–N_2_O values by 14.1 ± 1.5 mUr and δ^15^N^β^–N_2_O values by 4.5 ± 2.0 mUr (*n* = 3; Fig. 3e,f). Under the same conditions, CO_2_ mixing ratios increased by 4.20 % and 4.33 % (*n* = 2; Fig. 3b), respectively, while O_2_ decreased by 0.34 % and 3.83 % (*n* = 2; Fig. 3c).

**Fig. 3:**
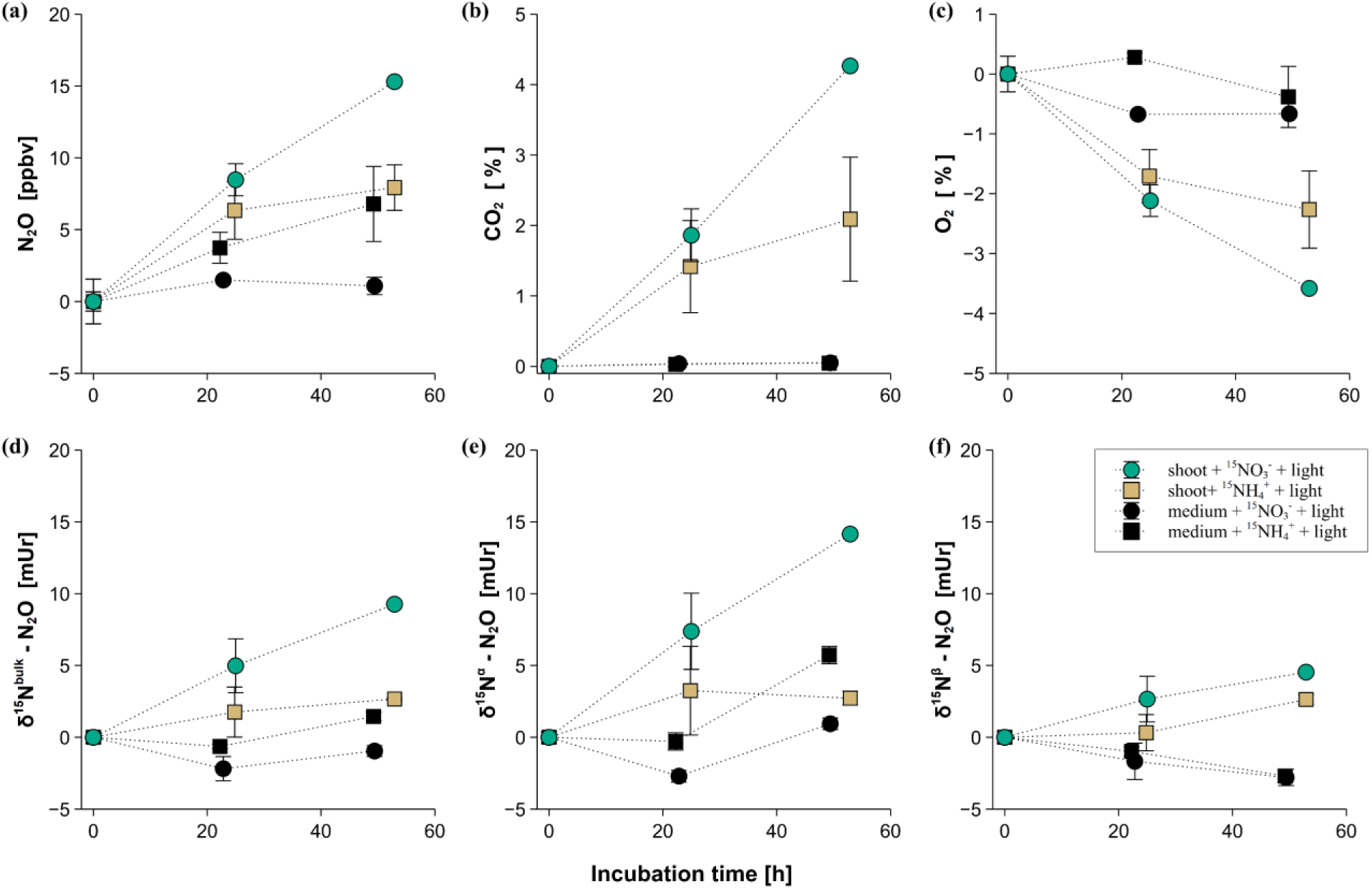
Relative changes of (a) N_2_O, (b) CO_2_ and (c) O_2_ mixing ratios and (d) δ^15^N^bulk^-N_2_O, (e) δ^15^N^α^-N_2_O and (f) δ^15^N^β^-N_2_O values during the incubation of chicory shoots under light exposure with supplementation of ^15^NO_3_^−^ and ^15^NH_4_^+^. Please note that the results are presented as changes relative to the measurements at the incubation start. Error bars represent the standard deviation (*n* = 3 or 4). Reported values reflect absolute changes in the respective glass flasks and were not normalized to biomass or its variation over time.

When shoots were supplied with ^15^NH_4_⁺ under light exposure, the N_2_O mixing ratio increased by 7.9 ± 1.6 ppbv (*n* = 3; Fig. 3a). Shifts towards more positive δ^15^N^bulk^–N_2_O values were more modest (+2.7 ± 0.5 mUr; *n* = 3; Fig. 3d) and increased by 2.7 ± 0.5 mUr for δ^15^N^α^–N_2_O and 2.6 ± 0.3 mUr for δ^15^N^β^–N_2_O (*n* = 3; Fig. 3e,f). CO_2_ mixing ratios increased by 2.09 ± 0.88 % and O_2_ decreased by 2.26 ± 0.64 % (*n* = 3; Fig. 3b,c).

Controls (‘medium + ^15^NO_3_⁻ + light’ and ‘medium + ^15^NH_4_⁺ + light’; *n* = 3) showed only minor increase in N_2_O mixing ratios of 1.1 ± 0.6 ppbv and 6.8 ± 2.6 ppbv, respectively (Fig. 3a). Isotopic shifts in these controls were negligible for δ^15^N^bulk^–N_2_O values (1.0 ± 0.4 mUr and +1.5 ± 0.3 mUr), while δ^15^N^α^–N_2_O/δ^15^N^β^–N_2_O values changed by <3 mUr (*n* = 2; Fig. 3d–f). CO_2_ and O_2_ mixing ratios remained within ± 0.4 % of initial values.

### Rates of N_2_O, CO_2_ production and O_2_ uptake

Root cultures produced N_2_O with rates of 15.1 ± 4.1 pmol h^-1^ g^-1^ (*n* = 3, *p* = 0.11) and 20.0 ± 5.1 pmol h^-1^ g^-1^ (*n* = 3, *p* = 0.21) for NO_3_^-^ and NH_4_^+^ supplementation, respectively. Shoots incubated under dark conditions exhibited the highest N_2_O production rates when supplemented with either ^15^NO_3_^−^ (23.2 ± 3.7 pmol h^-1^ g^-1^, *n* = 3) or ^15^NH_4_^+^ (27.1 ± 6.0 pmol h^-1^ g^-1^, *n* = 4), while the ‘shoot + dark only’ treatment, exhibited a production rate of 13.6 ± 2.8 pmol h^-1^ g^-1^ (*n* = 3, *p* < 0.05) (Fig. 4a). When shoots were incubated under light exposure, substantially lower N_2_O production rates were observed compared to those under dark conditions (*p* < 0.05). Specifically, the ‘shoot + ^15^NO_3_^−^ + light’ treatment yielded a production rate of 2.6 pmol h^-1^ g^-1^ (*n* = 2), while the ‘shoot + ^15^NH_4_^+^ + light’ treatment even showed a small net uptake of N_2_O, with a rate of –1.3 ± 0.8 pmol h^-1^ g^-1^ (*n* = 3).

**Fig. 4:**
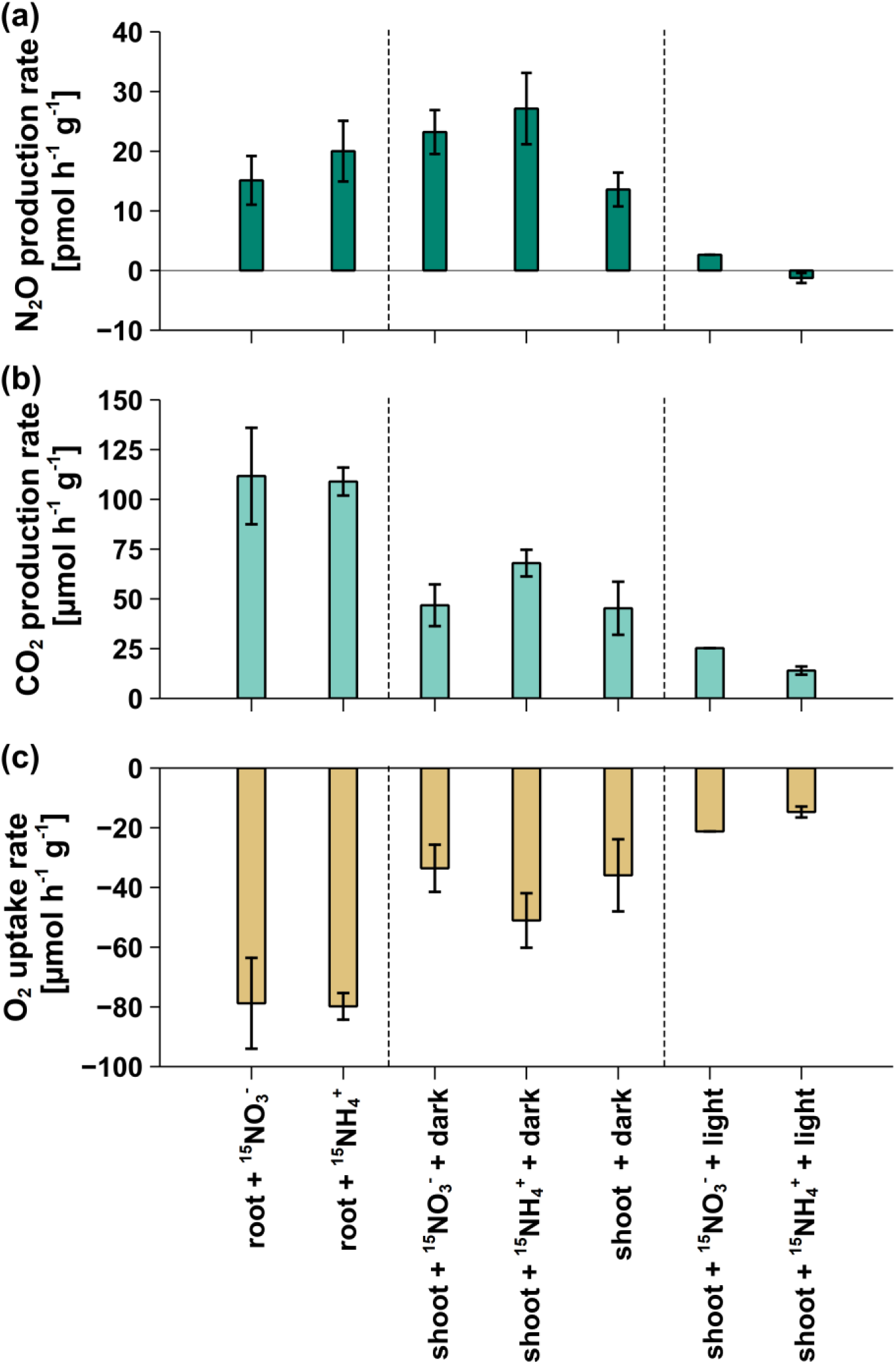
Production rates of (a) N_2_O, (b) CO_2_ and (c) uptake rates of O_2_ during incubation experiments of chicory root and shoot cultures. The results were normalized for plant dry weight (h^-1^ g^-1^). Error bars reflect propagated standard deviations (*n* = 3 or 4).

CO_2_ production rates were highest in ‘root + ^15^NO_3_^−^’ and ‘root + ^15^NH_4_^+^’ treatments, with rates of 111.7 ± 24.2 µmol h^-1^ g^-1^ (*n* = 3) and 109.0 ± 7.0 µmol h^-1^ g^-1^ (*n* = 3), respectively. Shoots incubated in the dark showed CO_2_ production rates of 46.8 ± 10.5 µmol h^-1^ g^-1^ (*n* = 3) for the ‘shoot + ^15^NO_3_^−^ + dark’ treatment, 68.0 ± 6.7 µmol h^-1^ g^-1^ (*n* = 4) for ‘shoot + ^15^NH_4_^+^ + dark’, and 45.3 ± 13.3 µmol h^-1^ g^-1^ (*n* = 3) for ‘shoot + dark only’. The lowest CO_2_ production rates were found in light-incubated shoots treated with ^15^NO_3_^−^ (25.3 µmol h^-1^ g^-1^, *n* = 2) and ^15^NH_4_^+^ (14.0 ± 2.1 µmol h^-1^ g^-1^, *n* = 3).

Root cultures also exhibited the highest O_2_ consumption rates, with values of 78.8 ± 15.2 µmol h^-1^ g^-1^ (*n* = 3) and 79.8 ± 4.5 µmol h^-1^ g^-1^ (*n* = 3) for ^15^NO_3_^−^ and ^15^NH_4_^+^ treatments, respectively. Shoots incubated under dark conditions showed lower O_2_ uptake rates compared to roots, with rates of 33.6 ± 7.9 µmol h^-1^ g^-1^ (*n* = 3) for ‘shoot + ^15^NO_3_^−^ + dark’, 51.1 ± 9.1 µmol h^-1^ g^-1^ (*n* = 4) for ‘shoot + ^15^NH_4_^+^ + dark’, and 35.9 ± 12.1 µmol h^-1^ g^-1^ (*n* = 3) for the ‘shoot + dark only’ treatment. Light-incubated shoots showed the lowest O_2_ consumption rates with 21.2 µmol h^-1^ g^-1^ (*n* = 2) when ^15^NO_3_^−^ and 14.7 ± 1.8 µmol h^-1^ g^-1^ (*n* = 3) when ^15^NH_4_^+^ was added.

### Change rates of δ^15^N-N_2_O

Root cultures supplemented with ^15^NO_3_^−^ showed the highest δ^15^N-change rates compared to all other root and shoot treatments (*p* < 0.02). Chicory roots exhibited clear differences in δ^15^N-N_2_O change rates, with δ^15^N^bulk^-N_2_O change rates of 1.53 ± 0.22 mUr h^-1^ g^-1^ (*n* = 3) and 0.27 ± 0.03 mUr h^-1^ g^-1^ (*n* = 3) following the addition of ^15^NO_3_^−^ and ^15^NH_4_^+^, respectively (*p* = 0.01). Dark-incubated shoots treated with ^15^NO_3_^−^ and ^15^NH_4_^+^ showed δ^15^N^bulk^-N_2_O change rates of 0.37 ± 0.06 mUr h^-1^ g^-1^ (*n* = 3) and 0.35 ± 0.06 mUr h^-1^ g^-1^ (*n* = 4), respectively. Dark-incubated shoots without ^15^NO_3_^−^/^15^NH_4_^+^ supplementation showed a small increase in δ^15^N over time, with a rate of 0.24 ± 0.10 mUr h^-1^ g^-1^ (*n* = 3). Among light-incubated shoots, the lowest δ^15^N^bulk^-N_2_O rates were observed, with values of 0.17 mUr h^-1^ g^-1^ (*n* = 2) for the treatment ‘shoot + ^15^NO_3_^−^ + light’ and 0.04 ± 0.01 mUr h^-1^ g^-1^ (*n* = 3) for ‘shoot + ^15^NH_4_^+^ + light’.

When observing the position-specific δ^15^N change rates root cultures treated with ^15^NO_3_^−^ consistently exhibited higher rates compared to those treated with ^15^NH_4_^+^ (*p* < 0.02 for both δ^15^N^α^-N_2_O and δ^15^N^β^-N_2_O change rates). Specifically, δ^15^N^α^-N_2_O change rates were 1.68 ± 0.24 mUr for ‘root + ^15^NO_3_^−^’ (*n* = 3) and 0.37 ± 0.02 mUr for ‘root + ^15^NH_4_^+^’ (*n* = 3), whereas δ^15^N^β^-N_2_O change rates were slightly lower, at 1.40 ± 0.20 mUr and 0.18 ± 0.04 mUr, respectively, for the same treatments. The same trend of higher δ^15^N^α^-N_2_O change rates compared to δ^15^N^β^-N_2_O was also observed across all incubated shoot treatments. However, δ^15^N^α^-N_2_O and δ^15^N^β^-N_2_O change rates among shoot treatments displayed overall less variability and no clear pattern ranging from 0.18 to 0.51 mUr h^-1^ g^-1^ and 0.17 to 0.27 mUr h^-1^ g^-1^, respectively. A marginal negative δ^15^N^α^-N_2_O change rate of –0.13 ± 0.07 mUr (*n* = 3) was observed in the ‘shoot + ^15^NH_4_^+^ + light’ treatment.

## Discussion

Our experiments with sterile chicory cultures, combining concentration measurement and ^15^N-stable isotope labelling, demonstrate that both roots and shoots can independently produce and emit N_2_O without microbial influence. The isotopic enrichment of ^15^N in N_2_O, particularly in treatments with ^15^N-labelled NO_3_^−^ in root cultures, confirmed that plants themselves are the source of N_2_O. Combined with sterile conditions, these findings strongly suggest plant processes, rather than microbial pathways, were responsible for N_2_O production in our experiments. This is the first study to demonstrate that different plant parts, such as roots and shoots, produce N_2_O independently, based on distinct N_2_O production rates and isotopic changes during experiments with ^15^N-labelled precursors.

Both root and shoot cultures clearly showed N_2_O formation in our experiments. Roots consistently showed higher overall N_2_O mixing ratios than shoots. However, when normalized to dry biomass, N_2_O production rates were similar between roots and shoots, with shoot production rates exceeding root rates in treatments with added NO_3_^−^ and NH_4_^+^. Up to date, only few studies have reported plant N_2_O production rates normalized to biomass. Our observed N_2_O production rates (−1 to 27 pmol g^-1^ h^-1^) fall within the range reported by Lenhart *et al*. (2019; 14–6200 pmol g^-1^ h^-1^) for plants grown under sterile conditions but are lower than those reported by Qin *et al*. (2024; 114–330 pmol g^-1^ h^-1^). The lower rates compared to Qin *et al*. (2024), may reflect several factors. First, the absence of endophytic bacterial N_2_O production in our study, which were identified as the potential main driver of plant N_2_O emissions by these authors. Second, the lack of microbial activity that could support reduction steps from NO_3_^−^ to NO_2_^−^or NO_2_^−^ to NO. It is important to note that not all reduction steps need to be microbially driven. N_2_O formation may result from a combination of microbially and non-microbially catalyzed steps. The lower rates in our sterile conditions may therefore reflect purely biochemical N_2_O formation without the contribution of microbes that can catalyze or regulate parts of the reduction chain. Finally, species-dependent differences in N_2_O production, as demonstrated by Lenhart *et al*. (2019) may further contribute to the observed differences.

In our experiments, root cultures showed clear differences in δ^15^N^bulk^, δ^15^N^α^, and δ^15^N^β^-N_2_O change rates compared to shoot cultures (Fig. 5). N_2_O isotopic values were substantially more enriched in ^15^N when ^15^N-labelled NO_3_^−^ was added compared to NH_4_^+^, highlighting that most likely only ^15^N-NO_3_^-^ was converted to N_2_O in roots while NH_4_^+^ was not. This suggests that likely only NO_3_^-^ but not NH_4_^+^ contributed to N_2_O production in chicory root aligning with a number of previous studies (e.g., Goshima *et al*., 1999 and Lenhart *et al*., 2019).

**Fig. 5:**
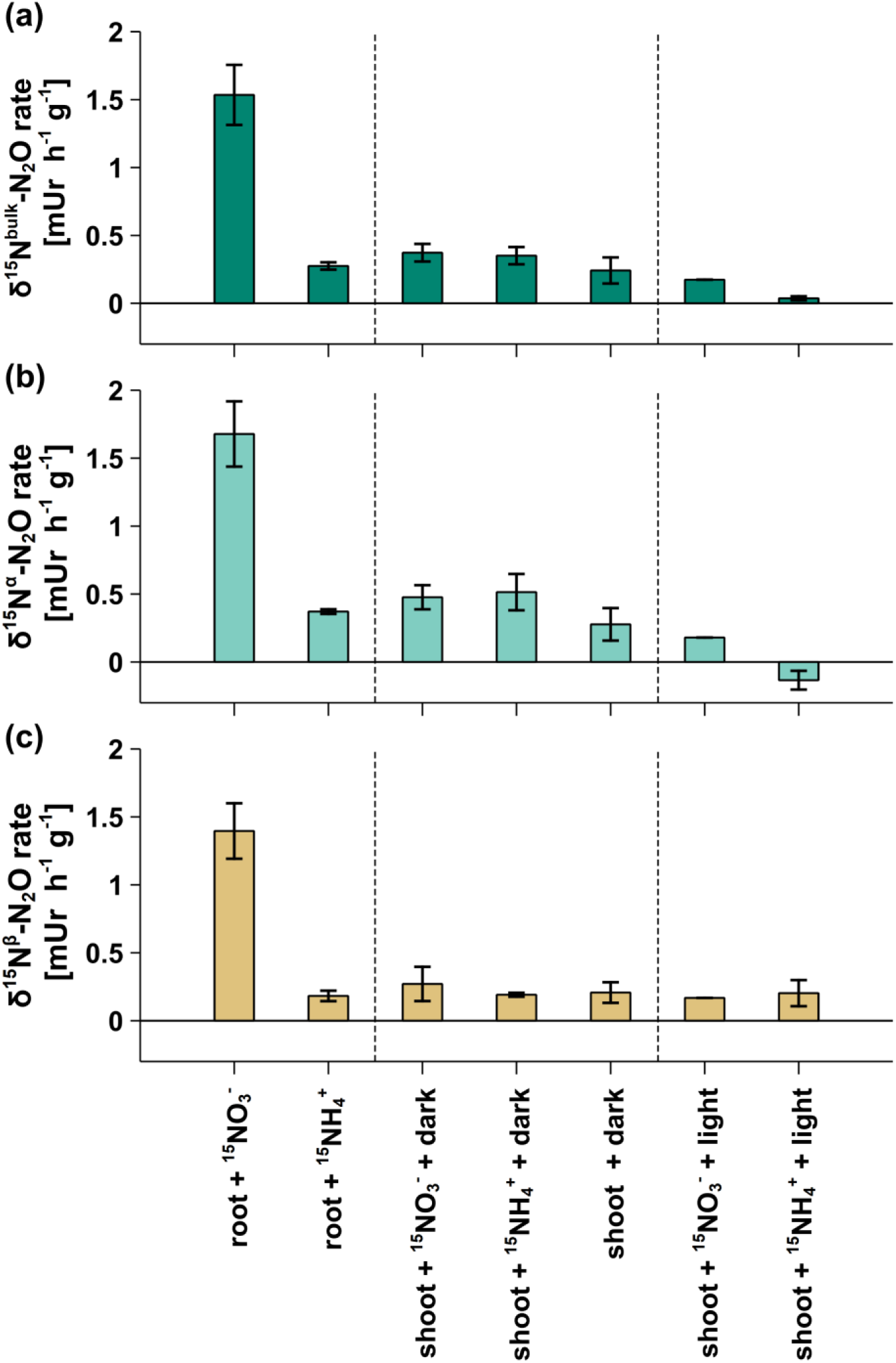
Isotopic change rates of (a) δ^15^N^bulk^, (b) δ^15^N^α^ and (c) δ^15^N^β^ in N_2_O of incubation experiments with chicory shoot and root cultures. Obtained rates were normalized to plant dry weight (h^-1^ g^-1^). Error bars reflect propagated standard deviations (*n* = 3 or 4).

Interestingly, the ‘root + ^15^N-NO_3_^−^’ treatment was the only treatment, where both δ^15^N^α^-N_2_O and δ^15^N^β^-N_2_O values increased in a similar amount, while for all shoot treatments this was not the case. Notably, this stable isotope observation clearly highlights that both nitrogen atoms from the N_2_O were likely of the same origin (the supplemented ^15^N-NO_3_^-^), since regardless of the position of N^α or β^-N_2_O, ^15^N-was enriched. Hence, N_2_O formation via the NO-pathway, as described by Timilsina *et al*. (2020) was likely responsible for the observed N_2_O formation in this treatment. In this context, root-associated N_2_O formation from nitrate (NO_3_⁻) appears to be linked to higher oxygen consumption by the roots, as indicated by elevated CO_2_ mixing ratios (Figs. 1–3). Reduced O_2_ levels in the root incubation flasks likely decreased internal root O_2_, which can stimulate (NO_3_⁻) reduction to (NO_2_⁻), (NO), and (N_2_O) through mitochondrial cytochrome c oxidase activity (Timilsina *et al*., 2020). Under low-oxygen (hypoxic) conditions, the reduction of NO_2_⁻ to NH_4_^+^ by nitrite reductase is inhibited. This favors NO accumulation and its subsequent conversion to N_2_O via the phytoglobin–NO cycle. Such a pathway may help plants reduce nitrosative stress by removing toxic NO, while at the same time representing an important global source of biogenic N_2_O. Timilsina *et al*. (2020, 2022a) further suggested that mitochondria play a central role in plant-derived N_2_O formation, which is consistent with the observed correlation between N_2_O production and respiration rates in our study.

When comparing ^15^N enrichment in N_2_O among treatments, the ‘shoot + dark’ treatment showed the strongest ^15^N enrichment in δ^15^N^bulk^-N_2_O and δ^15^N^α^-N_2_O change rates, whereas δ^15^N^β^-N_2_O showed little variation. This pattern was particularly clear in dark-incubated shoots, where δ^15^N^β^-N_2_O change rates remained consistent across all treatments, even in the absence of ^15^N-labelled substrates.

In shoots incubated in the dark and supplied with ^15^N-NO_3_⁻ or ^15^N-NH_4_^+^, N_2_O production rates increased compared to the ‘shoot + dark’ treatment (Fig. 4). However, the observed isotope shifts could not be conclusively attributed to the addition of ^15^N-labelled substrates, since the ^15^N-change rates were similar to those in the ‘shoot + dark’ treatment without added ^15^N-NO_3_⁻ or ^15^N-NH_4_^+^. This suggests that N_2_O formed under dark conditions alone may be responsible for the observed isotope patterns rather than direct conversion of the labelled nitrogen sources.

Similar to the dark-incubated root experiments, darkness in shoots likely enhances respiration, reducing internal O_2_ levels and creating more favorable conditions for NO and N_2_O formation. This balance between photosynthetic O_2_ release and mitochondrial O_2_ consumption may be a critical factor determining whether N_2_O accumulates or is further metabolized. Under low-O_2_ (hypoxic or anoxic) conditions, nitrate reductase shifts its activity from NO_3_⁻ reduction toward NO_2_⁻-dependent NO formation due to restricted electron transport in respiration. In contrast, under high-O_2_ conditions, NO formation is suppressed because aerobic respiration dominates and NO-scavenging pathways are more active.

The influence of internal O_2_ levels on N_2_O formation pathways was also evident in shoots incubated under light conditions. In these treatments, ^15^N enrichment in N_2_O was low, suggesting that conversion of supplemented ^15^N-NO_3_⁻ or ^15^N-NH_4_^+^ to N_2_O occurred, if at all, only to a minor extent. Shoots supplied with ^15^N-NH_4_^+^ and ^15^N-NO_3_⁻ under light showed only small differences in isotope values between treatments. As discussed above, these small variations most likely reflect the intrinsic isotope signature of N_2_O produced by the shoots themselves, rather than incorporation of ^15^N from the supplied substrates. Due to technical issues, the ‘shoot + light’ control treatment could not be included in this comparison.

Interestingly, under light conditions, N_2_O production was reduced by 88% in the ‘NO_3_^−^ + light’ treatment, while the ‘NH_4_^+^ + light’ treatment resulted in net N_2_O uptake. This suggests that N_2_O formation is strongly inhibited under light or that part of the produced N_2_O in the plant tissue is subsequently reduced to N_2_ (Timilsina *et al*., 2020, 2022a). This observation aligns with Schützenmeister et al. (2020), who observed up to a 63% reduction in N_2_O production in ash and beech saplings during photosynthesis. Photosynthesis likely increases internal O_2_, reducing the low O_2_ conditions required for alternative reduction pathways (NO_2_^−^ → NO → N_2_O). The reduction in N_2_O emissions under light suggests that plant metabolic activity may influence denitrification. This may be linked to cytochrome c oxidase in plant mitochondria, which shares similarities with bacterial reductases and could facilitate N_2_O reduction to N_2_ (Timilsina et al., 2020). Evidence supporting this phenomenon includes N_2_ emissions from wheat crops supplied with labeled nitrite (NO_2_^−^), indicating mitochondrial involvement in N_2_O reduction (Vanecko & Varner, 1955). The negative correlation between N_2_O consumption and CO_2_ respiration in plants and lichens further supports this hypothesis (Timilsina et al., 2020). Light further enhances mitochondrial activity and O_2_ availability, which may suppress N_2_O accumulation by promoting efficient electron transport and respiration. Under light conditions, NO_3_^−^ assimilation is efficient, preventing NO_2_^−^ accumulation, a key N_2_O precursor (Timilsina et al., 2020). This efficiency is driven by nitrite reductase, which reduces NO_2_^−^ to NH_4_^+^ thereby minimizing NO_2_^−^ availability for N_2_O formation (Oliveira et al., 2013; Chamizo-Ampudia et al., 2017). This mechanistic link between light-dependent nitrogen assimilation and reduced N_2_O production highlights the importance of photosynthetic activity in regulating plant-mediated N_2_O emissions. However, further studies are needed to clarify the enzymatic mechanisms behind this process.

In contrast to other studies, such as Lenhart *et al*. (2019), which found no effects of light on N_2_O emissions by plants, Bruhn *et al*. (2014) even reported increased N_2_O production rates when plants were exposed to light. However, in the latter study, Bruhn *et al*. (2014) proposed a mechanism of N_2_O formation in plants involving ultraviolet (UV) light-induced N_2_O release from epicuticular waxes. This abiotic process is primarily observed under high temperatures and strong UV exposure. Since our experimental setup did not exposure to UV light, this pathway is unlikely to be relevant in our study. Nonetheless, it should be considered under field conditions.

Our observations align with recent studies suggesting that under certain environmental conditions, plants, such as European beech trees, can act as N_2_O sinks, absorbing N_2_O from the atmosphere. However, the underlying pathways of this phenomenon remain unclear and may be linked to the previously discussed reduction of N_2_O to N_2_ (Machacova *et al*., 2024)

Our results show and confirm that NO_3_^−^ is a key precursor of plant-mediated N_2_O formation especially in root cultures, as indicated by increased δ^15^N-N_2_O values when ^15^N-NO_3_^−^ was supplemented, consistent with previous studies (Goshima *et al*., 1999; Hakata *et al*., 2003; Lenhart *et al*., 2019).

Importantly, all controls with medium (with or without ^15^N-labelled NO_3_^−^ or NH_4_^+^) showed negligible changes in δ^15^N^bulk, α-, or β^-N_2_O values, however there was a slight increase in δ^15^N^bulk^-N_2_O and δ^15^N^α^-N_2_O values observed for the ‘medium + ^15^NO_3_^−^ + dark’ control (Figs. 1-3). While the exact mechanism for this observed increase in δ^15^N-N_2_O values is unclear, it may indicate a minor abiotic pathway for N_2_O formation from ^15^NO_3_^−^. However, the observed increase was much smaller compared to incubations containing roots or shoots, suggesting that plant tissue is the primary driver of ^15^N-enrichment and N_2_O production.

These different isotopic patterns from our experiments suggest distinct N_2_O formation mechanisms in roots and clearly indicate that N_2_O formation in roots is linked to NO as a precursor, since the ^15^N from NO_3_^-^ was found in both the ^15^N^α^ and ^15^N^β^ positions of N_2_O. This observation underscores the importance of stable isotope techniques for identifying N_2_O formation pathways. Such approaches are essential to distinguish between root-, shoot-, and soil-derived N_2_O, thereby enabling more accurate characterization of plant-mediated N_2_O emissions in future studies. Our isotopic results provide new insights into N_2_O formation, with significant implications for assessing plant part-derived N_2_O dynamics, even though technical issues prevented direct comparison between shoot-only cultures under dark and light conditions. Despite this, significant differences in δ^15^N^bulk^ and δ^15^N^α^-N_2_O change rates across treatments suggest distinct N_2_O production mechanisms in plant parts, creating a basis for future investigations.

Our study, in agreement with a number of recent studies, thus challenges the long-held belief that plants only act as passive conduits for N_2_O, transporting it from soil to leaves or stems through xylem or bacterial activity in the phyllosphere (Bowatte *et al*., 2014, 2015; Machacova *et al*., 2019; Qin *et al*., 2024). Moreover, our isotopic labelling experiments provide direct evidence that plants are capable of producing N_2_O under sterile conditions, without any microbial influence, thereby offering new mechanistic insights beyond previous studies. Since our cultures were grown and incubated under sterile conditions, microbial pathways are unlikely to explain N_2_O production, which aligns with accumulating evidence suggesting that plants can actively produce N_2_O (e.g., Lenhart *et al*., 2019; Timilsina *et al*., 2022a).

Figure 6 schematically illustrates the current understanding of N_2_O production and uptake processes in plants, as well as the newly identified processes discovered in this study. Most current studies focus on microbial activity as the primary source of N_2_O, with limited consideration of direct plant production. Distinguishing microbial from plant-based N_2_O formation pathways is crucial for refining N_2_O cycling models and understanding their climatic impact. Our findings suggest that plants not only produce N_2_O, but that factors such as light can strongly influence N_2_O formation in roots and shoots or even lead to N_2_O uptake. The interplay of these factors requires further investigation. Overall, our results highlight that N_2_O dynamics in the soil–plant–atmosphere continuum are more complex than previously recognized. In addition to soil-derived N_2_O transported through plants, N_2_O produced in roots and shoots may mix as it moves upward, with some of it potentially being reduced within plant cells. This could result in either net N_2_O emission or uptake, depending on plant species, soil properties, and environmental conditions. Furthermore, the phyllosphere may host both N_2_O-producing and N_2_O-consuming bacteria, adding another layer of N_2_O dynamics that can influence overall emissions or uptake. This improved understanding of plant-mediated N_2_O processes has important implications for modeling N_2_O fluxes in the soil–plant–atmosphere system and should be incorporated into future vegetation-based N_2_O emission models, which currently do not account for these pathways (e.g., Lenhart *et al*., 2019; Qin *et al*., 2024).

**Fig. 6:**
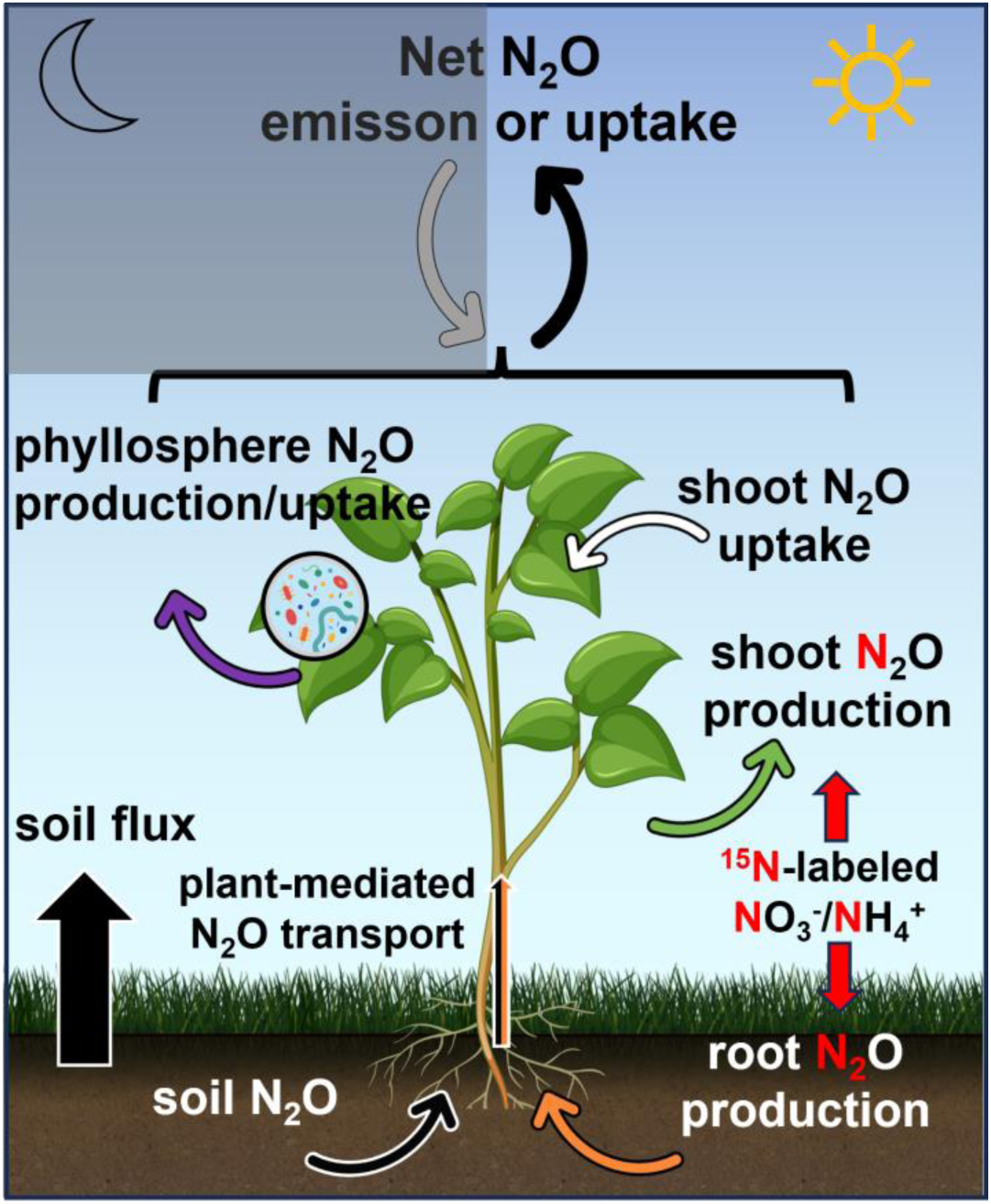
Schematic illustration of different processes for N_2_O production and uptake in soils and plants, illustrating the complex interaction of these processes leading to net N_2_O emissions or uptake by plants.

Building on the findings of this study, future research should aim to disentangle plant-derived N_2_O formation from microbial contributions. Key priorities include: (1) distinguishing root- and shoot-associated N_2_O pathways and their interactions with microbial processes, and (2) investigating N_2_O formation in shoots under dark conditions to separate plant-specific production from microbial effects. These aspects are critical for understanding how plants act as conduits or modifiers of N_2_O fluxes. Further studies should also (3) examine how different levels of photosynthetically active radiation affect N_2_O production and uptake across plant tissues, potentially revealing threshold or dose-dependent effects, and (4) explore the absence of N_2_O production or consumption in light-exposed shoots. If consistent across species and conditions, light exposure could be leveraged to reduce N_2_O emissions from vegetated systems. Together, these insights are essential for accurately quantifying plant contributions to N_2_O budgets and for improving global upscaling of N_2_O fluxes from terrestrial ecosystems.

## Acknowledgments

This study was supported by the Heidelberg Center for the Environment (HCE; project ExJ.2.2 Az:0078.3.2.2) and the DFG (German Research Foundation; project GR 5871/2-1). We thank C. Walter for the cultivation of the chicory root and shoot cultures and K. Lenhart for valuable discussions and her advisory support. We are grateful to B. Knape and S. Rheinberger for technical support.

## Author Contribution

**Moritz Schroll:** Conceptualization, Methodology, Investigation, Formal analysis, Data curation, Visualization, Writing - Original Draft, Writing - Review & Editing, Project administration. **Maurice Maas:** Methodology, Investigation, Formal analysis, Data curation, Visualization, Writing - Original Draft, Writing - Review & Editing. **Thomas Klintzsch:** Conceptualization, Writing - Review & Editing. **Steffen Greiner:** Conceptualization, Methodology, Resources, Writing - Review & Editing, Project administration, Supervision, Funding acquisition. **Frank Keppler:** Conceptualization, Methodology, Resources, Writing - Review & Editing, Project administration, Supervision, Funding acquisition.

## Data Availability

The experimental data will be stored at Heidelberg University internal database, which is publicly available (after publication) upon request.

## Notes

### Competing Interest Statement

The authors have declared no competing interest.

