## Supplement for "Insights into plant-part specific N_2_O production in roots and shoots of chicory (*C. intybus*) using stable isotope labelling"

The following Supporting Information is available for this article:

**Method S1** Underlying principles of cross-sensitivities during analysis of $N_2O$ mixing ratios and isotope values using cavity ring-down spectroscopy.

CRDS instruments, like the Picarro G5131-*i* used in this study, are generally susceptible to be affected by cross-sensitivities. This term refers to the influence of other gases (e.g. $O_2$, $CO_2$ or $CH_4$) on the analytical result of the target gas (e.g., Harris *et al.*, 2020), in this case, $N_2O$. Additionally to cross-sensitivities, the mixing ratio of the target gas itself may have potential impact on analytical results, too. To account for these analytical problems, correction factors need to be applied to the results obtained from analytical measurements. These factors are determined through experiments involving known mixing ratios of the respective gases and their impact on measurements of $N_2O$ mixing ratio and isotopic composition. For the present study, these corrections were specifically established for the used Picarro G5131-*i* device. While previous studies have derived correction factors for common gases affecting this type of instrument, the specific effects of cross-sensitivities are device-dependent, necessitating the determination of correction factors through independent experiments (Harris *et al.*, 2020). In this study, we derived correction factors using linear regression analysis for the influence of $O_2$, $CO_2$, and $CH_4$, as well as for different $N_2O$ mixing ratios, on the measurements obtained with the Picarro G5131-*i*. Consistent with prior research, we observed a statistically significant correlation between the mixing ratios of $O_2$, $CO_2$, and $CH_4$ and the measured $N_2O$ mixing ratio and isotopic values. The exact methodology and sequence of applying these correction factors to the raw data are detailed in Method S3.

**Method S2** Underlying principles of the effect of vials, septa and storage time on cavity ring-down based $N_2O$ analysis.

The selection of vials and septa represents, together with cross-sensitivities, a major source of analytical error in CRDS-based $N_2O$ analysis. In some samples, we detected $N_2O$ mixing ratios below atmospheric levels, which cannot be explained with $N_2O$ loss related to diffusion. This effect was predominantly observed in control group samples, thereby excluding biological $N_2O$ degradation as a potential cause. These findings suggest that interactions between air samples and the materials of the vials or the white silicone-PTFE septa (IVA-Analysentechnik / manufacturer) have the potential to influence the analytical results. Underlying mechanisms could be physical and chemical processes such as absorption, desorption, or chemical reactions between $N_2O$ and the material of vials and septa (Laughlin & Stevens, 2003; Rochette & Bertrand, 2003). The logical choice for storage vials would have been Exetainers (Labco

Limited, UK), as they were also used for gas chromatographic analysis of $O_2$, $CO_2$ and $CH_4$ during this study and show the best results in relation to storage, tightness and $N_2O$ loss (Glatzel & Well, 2007). They also showed good results in tests regarding $\delta^{15}$N-$N_2O$ measurements via IRMS (Laughlin & Stevens, 2003). However, in our own experiments Exetainers revealed a significant shift in $\delta^{15}$N-values even shortly after gas injecting into Exetainers, which renders them unsuitable for our specific analytical requirements. Possible reason could likely be outgassing of septa compounds interfering with CRDS measurements. To address this issue, we conducted experiments evaluating three different septa types: blue silicone-PTFE septa (Chromatographie-Zubehör Trott / manufactuer), white silicone-PTFE septa (IVA-Analysentechnik / manufacturer) and butyl-PTFE septa (IVA-Analysentechnik / manufacturer). These experiments involved filling vials with ambient air and monitoring the changes in measured $N_2O$ mixing ratios and $N_2O$ isotopic composition over time. The results indicate that all three of the used septa types affected $N_2O$ mixing ratios and $\delta^{15}$N-$N_2O$ values. This effect was the smallest for the butyl-PTFE septa (IVA-Analysentechnik / manufacturer), making them the choice for subsequent experiments. Still, correction factors were required to account for these effects, which got stronger with increasing storage time. Thus, we derived time-dependent correction factors via linear regression analysis for the white silicone-PTFE septa (IVA-Analysentechnik / manufacturer) used in the earlier experiments and for the butyl-PTFE septa (IVA-Analysentechnik / manufacturer) we used in later experiments of this study. These correction factors have a temporal component to account for the different storage time of samples. The exact methodology and sequence of applying these correction factors to the raw data are detailed in Fig. S1 and Method S3.

**Fig. S1** Flow chart of steps from raw results to corrected results.

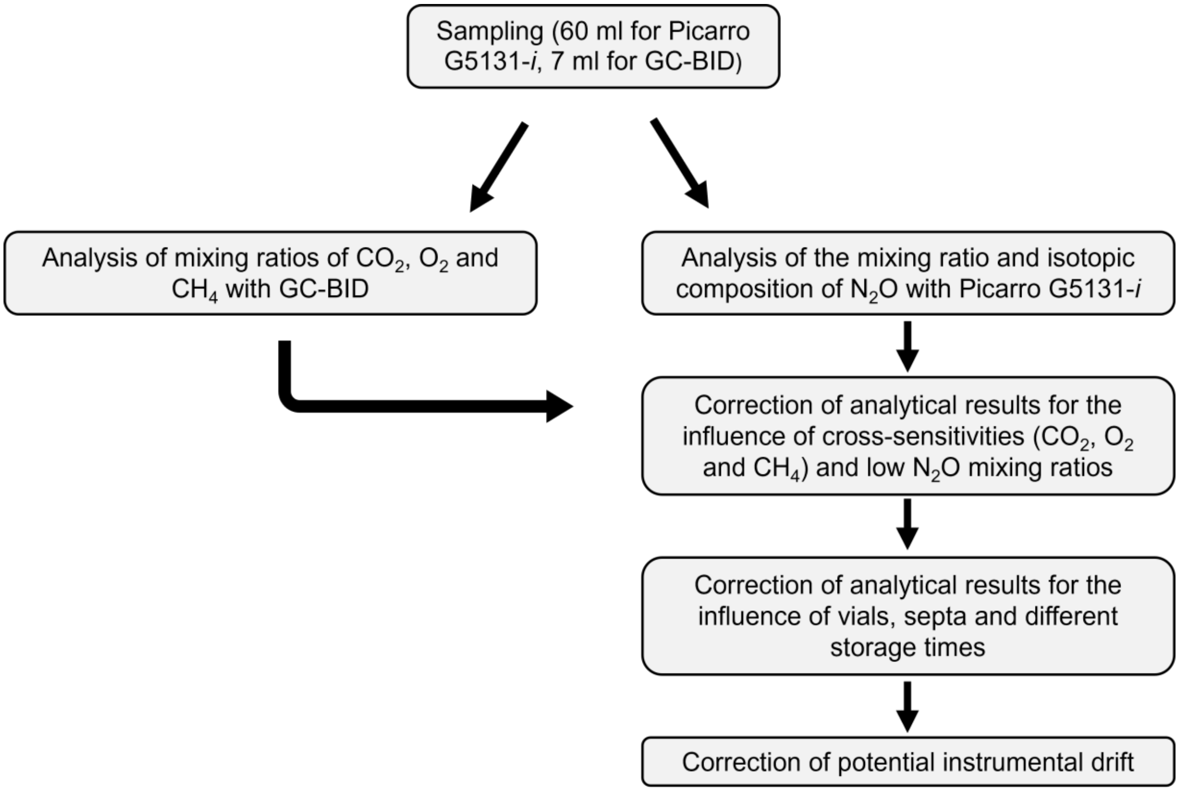

**Method S3** Processing and correction of raw data.

As outlined above, cross-sensitivities with other gases and low $N_2O$ mixing ratios, as well as septa, vials, and storage, influence the measurements of $N_2O$ mixing ratios and isotopic composition using the Picarro G5131-*i*. To minimize these effects, corrections were applied to the measurement results. Fig. S1 illustrates the workflow and the sequential steps of the corrections as applied during this study. The subsequent chapters provide a more detailed explanation of each correction step.

**Analysis of $CH_4$ mixing ratios**

$CH_4$ mixing ratios were determined using the same method as for $CO_2$ and $O_2$ mixing ratios (see Material and Methods), with a GC-2010 Plus gas chromatograph (Shimadzu, Japan) coupled with a barrier discharge ionization detector (BID-2010 Plus, Shimadzu, Japan) being used. (Setup: packed stainless-steel column (length: 2 m, inner diameter: 0.53 mm), stationary phase: carbon molecular sieve (ShinCarbon ST, 80/100 Mesh; Supelco, USA), carrier gas:  High-purity helium (grade 6.0, Air Liquide, France) at a constant flow rate of 5 mL $min^{-1}$).
The sample injection was performed using an autosampler (AOC-20i, Shimadzu, Japan). As part of quality control, one reference standard was measured after every six to nine single measurements. Calibration was performed with certified standard gas mixtures (CRYSTAL Gas Mixture, Air Liquide, France). Methane quantification was based on analyses of multiple reference standards ($CH_4$: 50–25,000 ppmv), measured in triplicates.

**Correction for cross-sensitivities**

The raw isotopic data were corrected for cross-sensitivities of $O_2$, $CO_2$, and $CH_4$. As an example, the correction of the $\delta^{15}N^{bulk}$ value for the cross-sensitivity with $O_2$ is illustrated using the following equations (Eqs. 1, 2 and 3). First, the difference ($\Delta O_2$) between the $O_2$ mixing ratio of the sample and that of the standard gas was determined:

$$\Delta O_2 = O_2 \text{ (sample)} - O_2 \text{ (standard gas)} \quad (1)$$

Subsequently, the change in $\delta^{15}N^{bulk}$ ($\Delta\delta^{15}N^{bulk}$) due to the cross-sensitivity with $O_2$ was calculated using the determined slope (m) of the regression line from experimental trials and $\Delta O_2$ determined from equation 1:

$$\Delta\delta^{15}N^{bulk} = m \times \Delta O_2 \quad (2)$$

The final corrected $\delta^{15}N^{bulk}$ value (denoted as $\delta^{15}N^{bulk}$ (C)) was obtained by adding $\Delta\delta^{15}N^{bulk}$ from the previous equation to the initial measured value ($\delta^{15}N^{bulk}$):

$$\delta^{15}N^{bulk} \text{ (C)} = \Delta\delta^{15}N^{bulk} + \delta^{15}N^{bulk} \quad (3)$$

Additionally to the correction for cross-sensitivities all measurement results with a $N_2O$ mixing ratio below 330 ppbv were corrected for the influence of a low $N_2O$ mixing ratio on isotopic measurements of $N_2O$. As an example, the correction of the $\delta^{15}N^{bulk}$ value is showed in the equations below (Eqs. 4 and 5). The difference in $\delta^{15}N^{bulk}$ ($\Delta\delta^{15}N^{bulk}$) caused by the low $N_2O$ mixing ratio was determined using the difference between 330 and the $N_2O$ mixing ratio and the slope (m) of the regression line from experimental trials (Figs S1-4):

$$\Delta\delta^{15}N^{bulk} = m \times (330 - N_2O \text{ [ppbv]}) \quad (4)$$

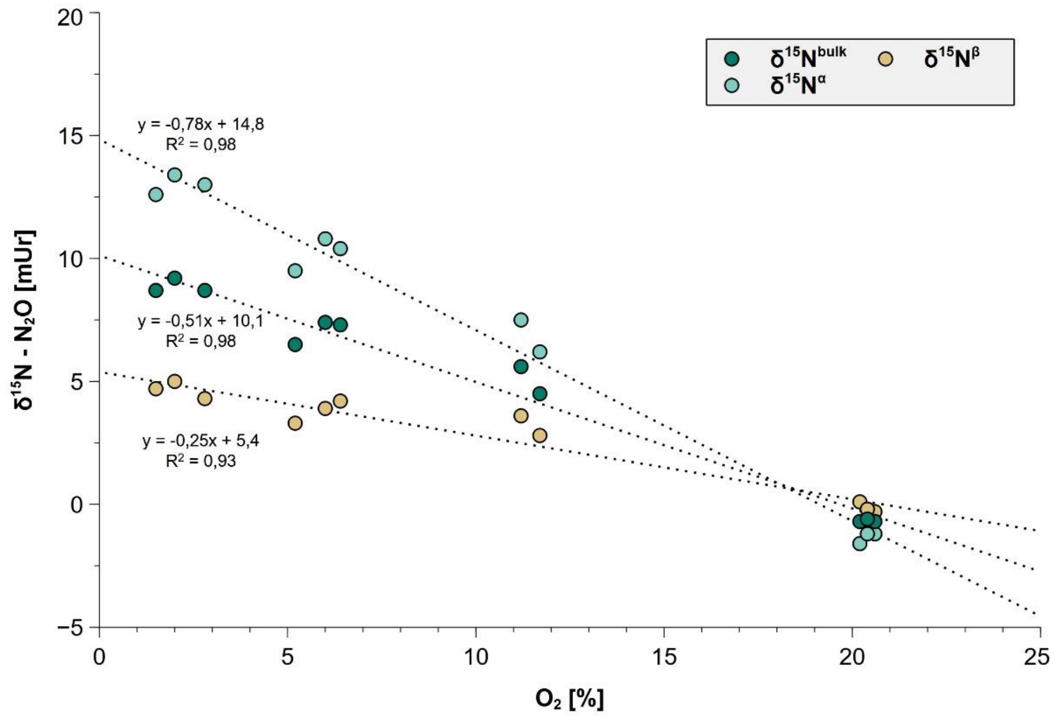


**Fig. S2** Change in $N_2O$ isotopic composition with varying $O_2$ mixing ratios. Correction factors were derived from linear regression, including $R^2$ values.

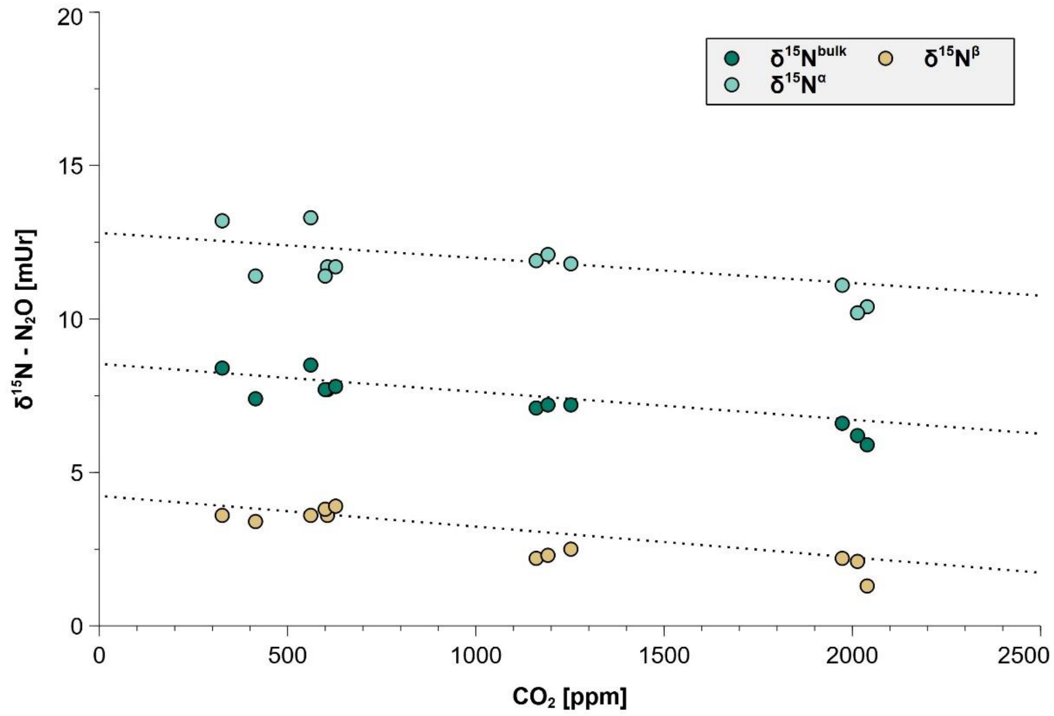


**Fig. S3** Change in $N_2O$ isotopic composition with varying $CO_2$ mixing ratios. Correction factors were derived from linear regression, including $R^2$ values.

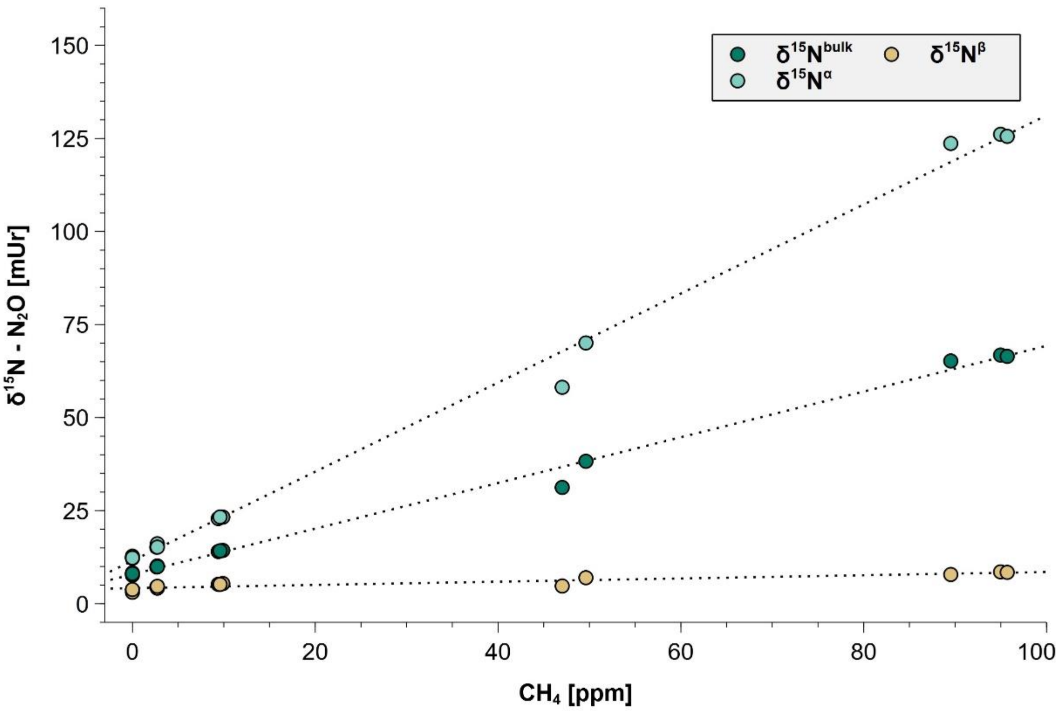


**Fig. S4** Change in $N_2O$ isotopic composition with varying $CH_4$ mixing ratios. Correction factors were derived from linear regression, including $R^2$ values.

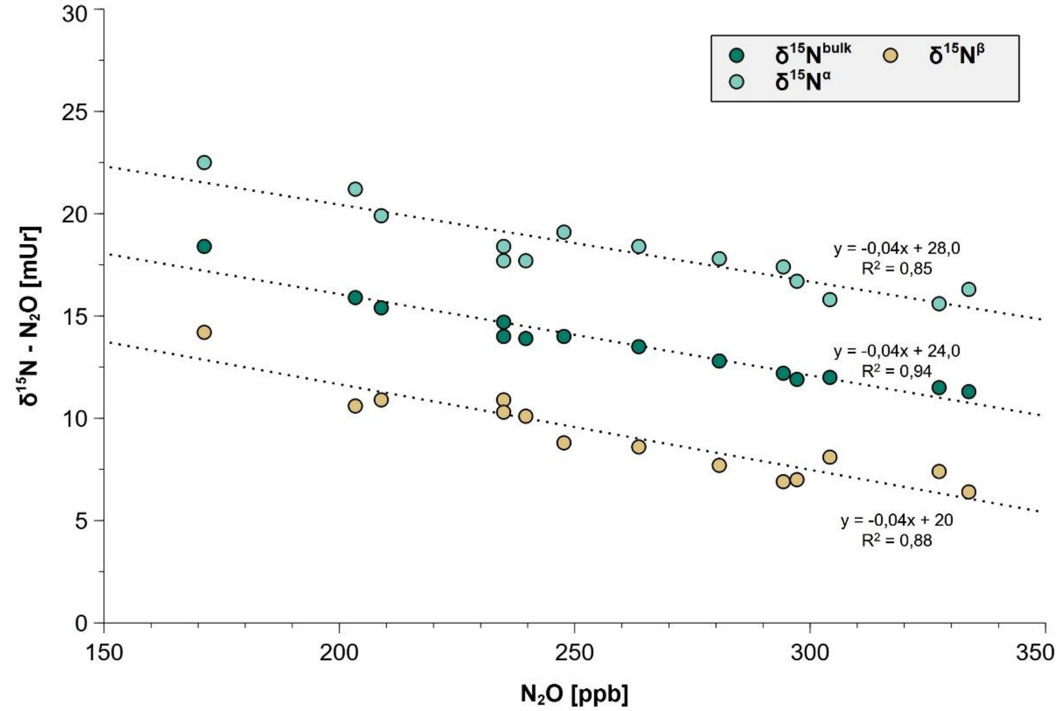


**Fig. S5** Change in $N_2O$ isotopic composition at $N_2O$ mixing ratios below 330 ppbv. Correction factors were derived from linear regression, including $R^2$ values.

The calculated value ($\Delta\delta^{15}N^{bulk}$) was then added to the initial measured value ($\delta^{15}N^{bulk}$) to determine the corrected $\delta^{15}N^{bulk}$ value ($\delta^{15}N^{bulk}$ (C):

$$\delta^{15}N^{bulk}\,(C) = \Delta\delta^{15}N^{bulk} + \delta^{15}N^{bulk} \tag{5}$$

**Correction for influence of septa, vials and storage**

In addition to the correction of the influence of cross-sensitivities from other gases and for the influence of $N_2O$ itself, the measurement results need to be corrected for the influence of septa and vials used for sample storage. The longer the period between sampling and measurement, the greater the effect of the septa on the measured values. The $\delta^{15}N^{bulk}$ value is used as an example for the correction (Eq. 6):

$$\delta^{15}N^{bulk}\,(C) = \delta^{15}N^{bulk} \times m \times t\,[h] \tag{6}$$

$\delta^{15}N^{bulk}$ refers to the initial measurement value, m is the slope of the regression line determined from the experimental trials, t is the time during which the sample remained in the vial with respective septa, and $\delta^{15}N^{bulk}$ (C) is the corrected value. Corrections factors of the respective septa can be found in Figs. S5-9:

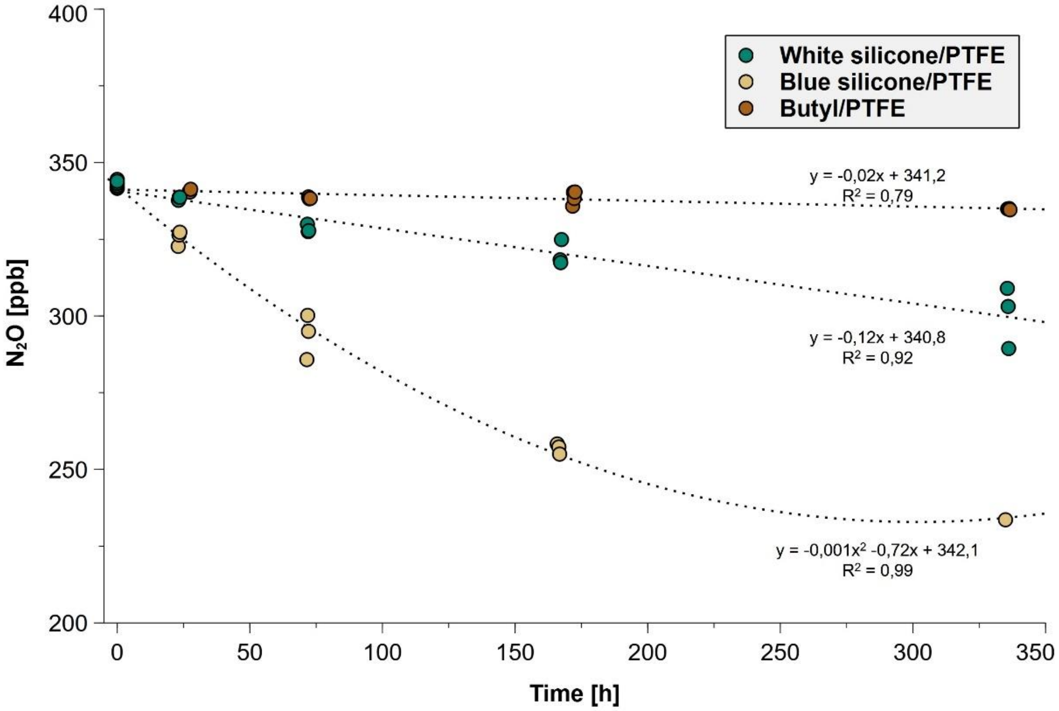


**Fig. S6** Change in measured $N_2O$ mixing ratios using different septa types (blue and white silicone/PTFE, butyl/PTFE) over time. Correction factors were derived from linear regression, including $R^2$ values.

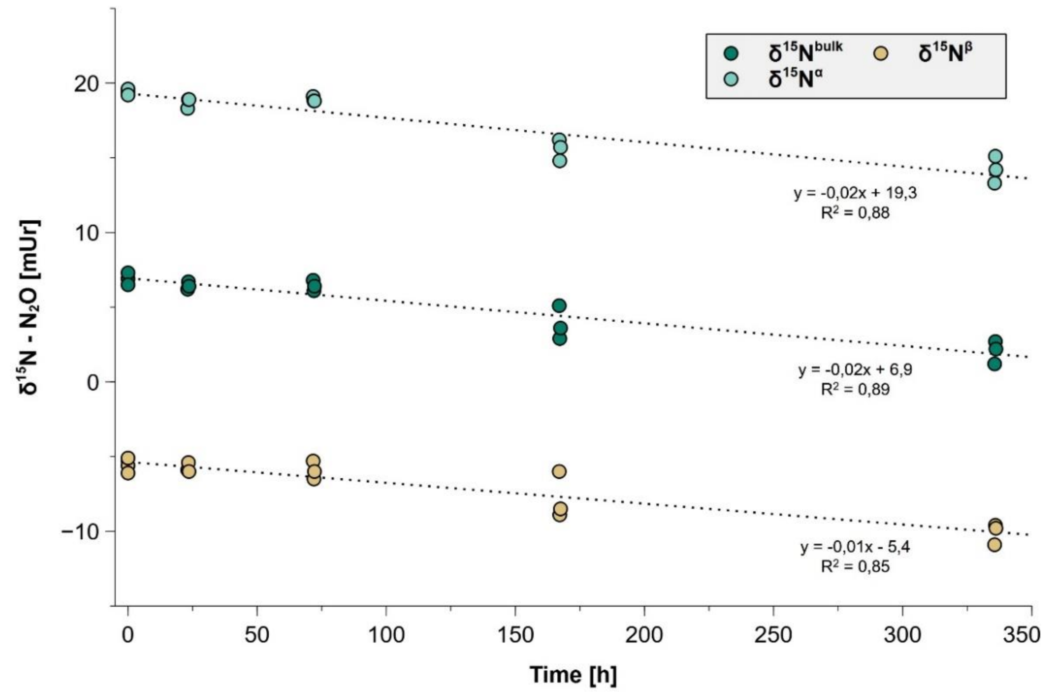


**Fig. S7** Change in $N_2O$ isotopic composition using white silicone/PTFE septa over time. Correction factors were derived from linear regression, including $R^2$ values.

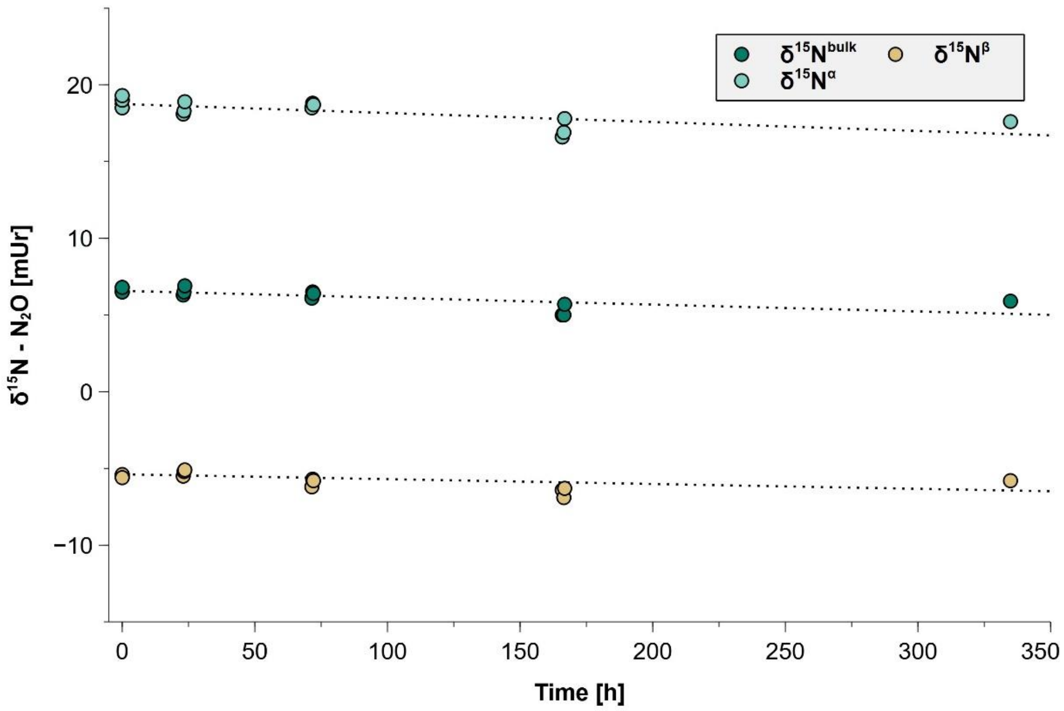


**Fig. S8** Change in $N_2O$ isotopic composition using blue silicone/PTFE septa over time. Correction factors were derived from linear regression, including $R^2$ values.

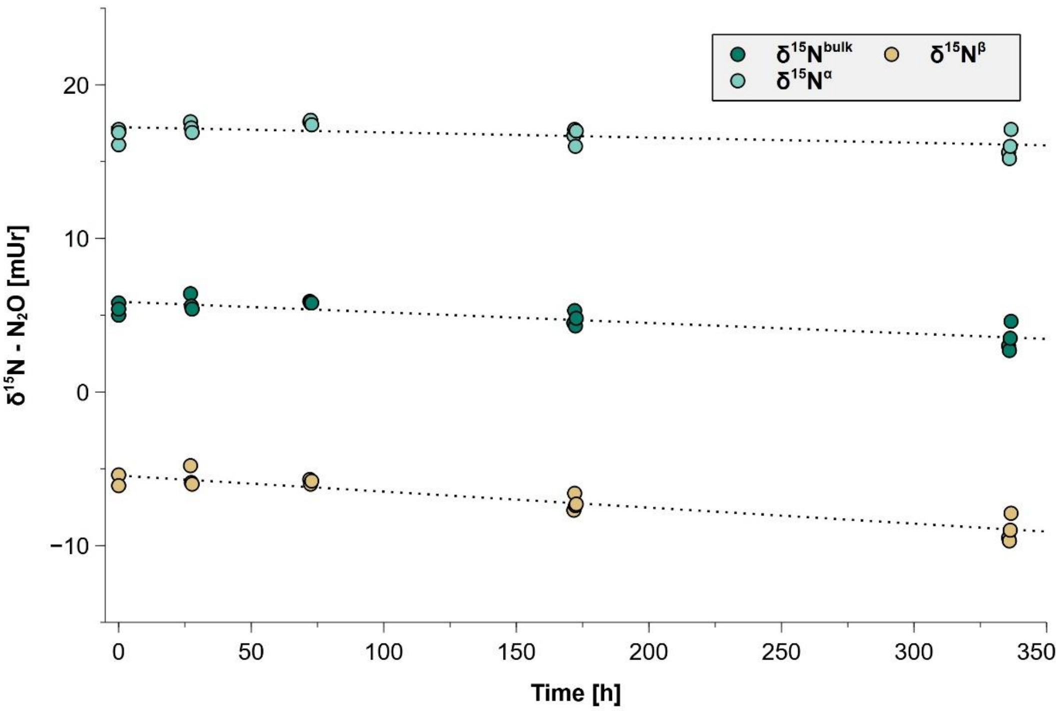


**Fig. S9** Change in $N_2O$ isotopic composition using butyl/PTFE septa over time. Correction factors were derived from linear regression, including $R^2$ values.

**Correction of potential instrument drift**

One in-house reference gas for $N_2O$ mixing ratios and one reference standard gas (RM 2; Mohn *et al.*, 2022) were analyzed at the beginning and end of each measurement day, allowing correction of potential instrumental drift by applying a linear two-point normalization following Paul *et al.* (2007).
